# Identification of a covert evolutionary pathway between two protein folds

**DOI:** 10.1101/2022.12.08.519646

**Authors:** Devlina Chakravarty, Shwetha Sreenivasan, Liskin Swint-Kruse, Lauren L. Porter

## Abstract

Although homologous protein sequences are expected to adopt similar structures, some amino acid substitutions can interconvert α-helices and β-sheets. Such fold switching may have occurred over evolutionary history, but supporting evidence has been limited by the: (1) abundance and diversity of sequenced genes, (2) quantity of experimentally determined protein structures, and (3) assumptions underlying the statistical methods used to infer homology. Here, we overcame these barriers by applying multiple statistical methods to a family of ~600,000 bacterial response regulator proteins. We found that their homologous DNA-binding subunits assume divergent structures: helix-turn-helix versus α-helix+β-sheet (winged helix). Phylogenetic analyses, ancestral sequence reconstruction, and AlphaFold2 models indicated that amino acid substitutions facilitated a switch from helix-turn-helix into winged helix. This structural transformation likely expanded DNA-binding specificity. Our approach uncovers an evolutionary pathway between two protein folds and provides methodology to identify secondary structure switching in other protein families.

## Introduction

Life is sustained by the chemical interactions and catalytic reactions of hundreds of millions of folded proteins. The structures and functions of these proteins are determined by their amino acid sequences^1^. As such, sequence changes have various functional effects, ranging from none to intermediate impairment to complete loss^2,3^, with biological outcomes ranging from no observable effect to debilitating disease^4–6^. While many historical studies indicate that amino acid variation can locally or globally unfold protein structure^7,8^, such changes typically do not remodel secondary structure, such as converting α-helices to β-sheets. These findings support the well-established observation that proteins with similar sequences have similar folds and execute similar functions. These similarities allow protein folds to be classified into families^9–11^ and underlie state-of-the-art protein structure prediction methods^12–14^.

Nevertheless, recent work shows that a subset of amino acid changes can switch secondary structure. This process has been called “evolutionary metamorphosis^15^” and “evolved fold switching^16^”. For instance, the most frequent non-Hodgkin-lymphoma-associated mutation observed in human mycocyte enhancer factor 2 (MEF2) switches a C-terminal α-helix to a β-strand, likely impeding MEF2 function^17^. Furthermore, numerous single mutations deactivate the cyanobacterial circadian clock by preventing a transformation that is critical for its normal function – the switch of its C-terminal subdomain from a βααβ fold to an αββα fold^18^. Finally, a single mutation can switch the fold and function of an engineered protein G variant from a 3-α-helix bundle that binds human serum albumin to an α/β-plait fold that binds immunoglobulins^19,20^.

These examples suggest that evolved fold switching of secondary structures may be one mechanism by which new protein folds originated in nature. If so, this evolutionary mechanism could ideally be identified by searching for homologous protein sequences with different experimentally determined structures (**Fig. 1a**). Similar approaches have successfully identified evolutionary relationships between protein fold families with conserved secondary structures but different tertiary arrangements^21,22^.

**Figure 1.**
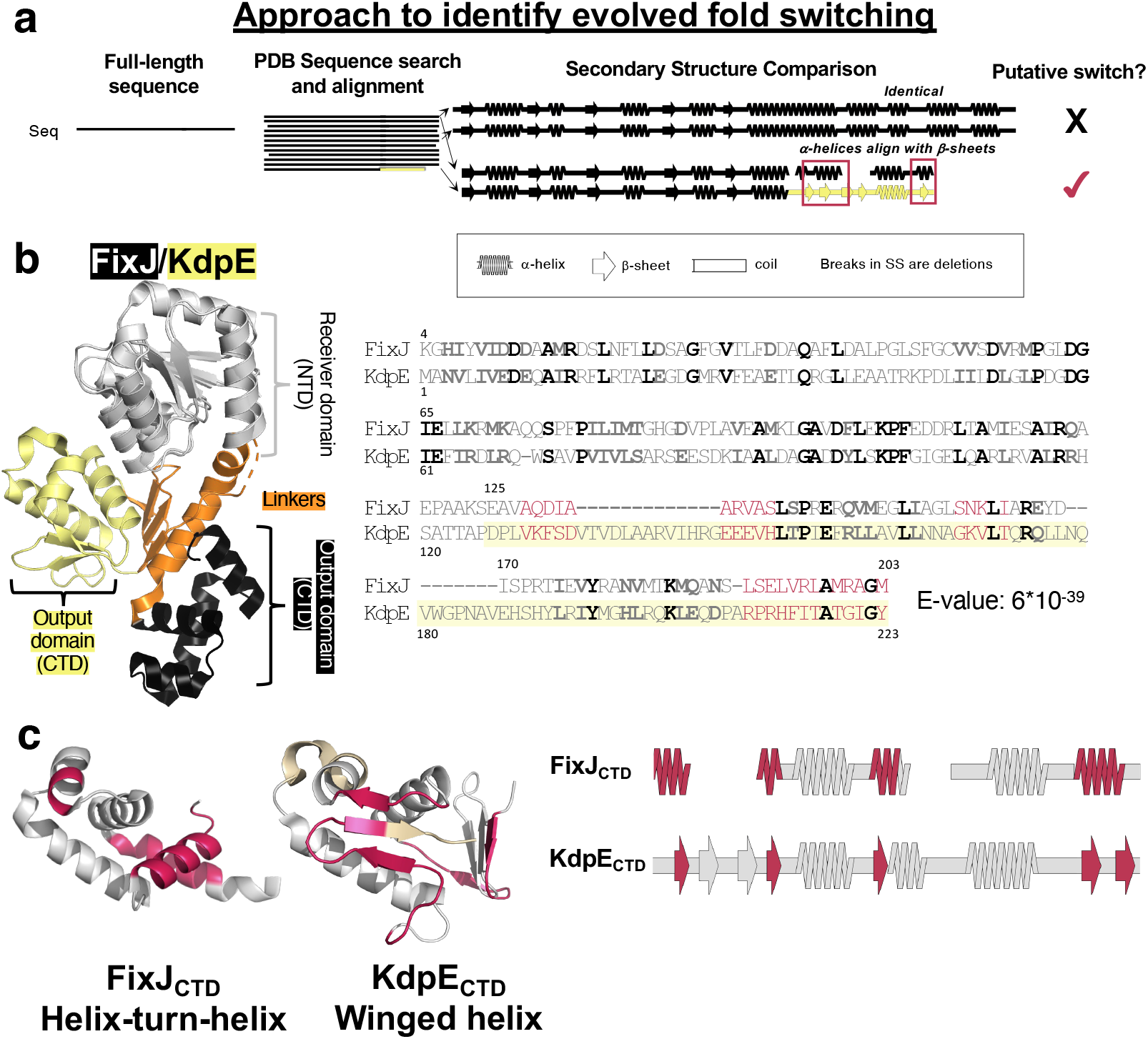
A combined sequence-structure search indicates that mutations may have switched some secondary structures between helix-turn-helix and winged helix proteins. (**a**). Querying the full sequence of FixJ (HTH_4_) against the PDB with one round of BLAST yielded a significant match with full-length KdpE (wH). Notably, in two regions, experimentally determined α-helices aligned with β-sheets. (**b**) A subsequent PSI-BLAST search confirmed a likely evolutionary relationship between the full-length FixJ and KdpE sequences; full-length structures are shown with conserved NTDs in gray, linkers in orange, HTH_4_ CTD in black, and wH CTD in yellow. The resulting PSI-BLAST alignment includes the NTD and CTD (starting where KdpE sequence is highlighted in yellow); bold amino acids are identical (black) or similar (gray), regions were α-helices align with β-strands are pink; gaps are denoted ‘-’. (**c**). Regions of three-dimensional structure (left) and secondary structure (right) where PSI-BLAST aligns α-helices in the HTH_4_ fold with sequences of β-strand in the wH fold (pink). Gray regions indicate conserved secondary and tertiary structure; beige regions in the wH correspond to its additional amino acids in the alignment, indicated as open spaces in the aligned secondary structure of FixJ (right).

However, observations of evolved secondary structure interconversion have been impeded by several technical barriers: 1) the limited abundance and diversity of sequenced genes, (2) the limited quantity of experimentally determined protein structures, and (3) the assumptions underlying the statistical methods used to infer homology. Indeed, all three limitations impacted the pioneering work of Cordes and colleagues, who identified a likely evolutionary relationship between the two distinctly folded transcription factors, P22 Cro and λ Cro^23–25^. Structurally, these two proteins share a 3-helical N-terminal core but have divergent C-terminal regions: P22 Cro’s C-terminal region folds into two α-helices, whereas λ Cro’s C-terminal assumes a β-hairpin. These differences could arise from an evolved fold-switching event. However, at the time of their study, this protein family comprised only 55 sequences and 5 solved structures, illustrating barriers (1) and (2). Illustrating barrier (3), whole-database PSI-BLAST searches did not suggest that P22 Cro and λ Cro were homologous, leading the authors to conclude^24^, “profile-based methods might be intrinsically ill suited… when wholesale structural change has occurred, since sequence conservation patterns will change in such a case.”

Here, we hypothesize that evolved fold switching can be observed with sufficient protein sequence and structure information. Since the aforementioned study was performed nearly 20 years ago, the number of available sequences in the RefSeq^26^ database has increased by three orders of magnitude, and the number of experimentally determined structures deposited in the Protein Data Bank (PDB) has increased by a factor of 7^27,28^. Further, more sensitive methods for inferring homology from sequences, specifically HMMER^29^ and HHblits^30^, have been developed.

Thus, to determine whether fold switching can now be observed in the evolutionary record, we searched for evidence among a large family of bacterial response regulators. We focused on a set of ~600,000 sequences and 76 unique, experimentally-determined structures. Each homolog constitutes one-half of a bacterial “two-component system” (TCS); the other half is a cognate sensor protein with an extracellular sensing domain and a histidine kinase domain^31^. When the extracellular domain of a sensor protein binds its triggering ligand, the histidine kinase domain phosphorylates its cognate response regulator protein at a conserved aspartate in the N-terminal receiver domain. In turn, this modification causes the response regulator’s C-terminal “output” domain to mount the organism’s response, such as altered transcription regulation^32^. This process allows bacteria to respond to their environments through chemotaxis^33^, antibiotic resistance^34^, oxygen sensing^35^, and more^36^.

Structurally, the response regulator proteins share a common N-terminal domain architecture. Structural differences between their C-terminal domains have been used to divide them into subfamilies^32,37^. Nearly 50% of the C-terminal domains fold into either helix-turn-helix (HTH) or winged helix (wH) DNA binding domains^32^; this ~50% corresponds to the ~600,000 mentioned previously. Both C-terminal domains comprise a core 3-helix bundle flanked by either (i) an N-terminal helical linker and a 4^th^ C-terminal helix (*e.g*., a tetrahelical HTH, or HTH_4_) or (ii) a four-stranded N-terminal β-sheet (here called a linker for ease of comparison) and a C-terminal β-hairpin (or “wing”, **Fig. 1b and c**). On average, response regulators with HTH_4_ output domains are ~30 residues shorter than their wH counterparts.

Common evolutionary descent of the response regulator HTH_4_ and wH domains was suggested previously.^38^ However, an evolutionary mechanism could not be detected, most likely due to the paucity of sequence and structure information available at the time of study. Thus, it has been unclear whether the differences in CTD secondary structures resulted from sequence insertions, complete or partial domain recombination, evolved fold switching, or some combination of the three.

In this work, we report strong statistical support for evolved fold switching of C-terminal secondary structure in HTH_4_ and wH domains and a putative evolutionary pathway between the two folds. First, we showed that the C-terminal α-helix of the HTH_4_ shares an evolutionary relationship with the β-sheet wing of the wH (**Figure 1**). This relationship was reinforced through multiple statistical analyses of phylogenetic relationships, ancestral sequence reconstruction with AlphaFold2 models, and functional analyses. All lines of evidence consistently point to an evolutionary trajectory by which an α-helix transformed into a β-sheet through stepwise mutation(s). Our results suggest how stepwise mutations can switch protein secondary structure and provide methodology to identify evolved fold switching in other protein families.

## Results

### Apparent homology between bacterial response regulators with HTH_4_ and wH CTDs

We previously used protein BLAST^39^ to identify pairs of protein sequences with high sequence identity (>70%) but divergent experimentally determined secondary structures^16,40^ (**Figure 1a**). In addition to sequence identities, BLAST searches report e-values, which quantify the statistical significance of a sequence match given the size of the database searched. Lower e-values indicate that a match is increasingly unlikely to arise by chance, allowing homology to be inferred. Since sequence pairs with <70% identity can also be homologous, we reapplied our original search strategy with a maximum e-value threshold of 1e-05. This is a conservative threshold since 5e-02 is often used to infer homology^39^, though some sequences with higher e-values are also homologous. Among our results, we identified a match between FixJ of *Bradyrhizobium japonicum* and KdpE from *Escherichia coli*.

Indeed, using the full-length sequence of FixJ_PDB_ to query the PDB retrieved the sequence of KdpE_PDB_ with an e-value of 1e-07. Both FixJ_PDB_ and KdpE_PDB_ are response regulators of bacterial two component systems. Their N-terminal domains (NTDs) showed high sequence and structural similarities (**Figure 1b**, left), whereas their linkers and DNA-binding C-terminal domains (CTDs) showed modest sequence similarities and striking differences in secondary structure: FixJ_PDB_’s CTD comprises a tetrahelical helix-turn-helix (HTH_4_) architecture, whereas KdpE_PDB_’s CTD comprises a winged helix (wH, **Figure 1**). The KdpE_PDB_ CTD is also 15 aa longer than that of FixJ_PDB_. Nonetheless, FixJ’s helical linker aligned partially with the four β-sheets of KdpE’s CTD. (For ease of comparison, we call both regions, “linkers”.) Furthermore, the C-terminal α-helix of FixJ_PDB_ aligns with the C-terminal β-hairpin of KdpE_PDB_’s CTD, also known as its “wing”.

In contrast, BLAST and PSI-BLAST searches of the PDB using the sequences of isolated CTDs from either FixJ or KdpE as queries only identified sequences from the same fold families (HTH_4_ or wH). Sequences encoding the alternative structure were not identified.

Two possibilities could explain these results. First, in the full-length sequences, the strong similarities of the NTD could erroneously give rise to the CTD alignment through “homologous overextension”, in which flanking, nonhomologous sequences are erroneously included in a local sequence alignment^41^. In this case, the distinctly folded CTDs would *not* share a common ancestor. Instead, genes encoding the separate CTDs likely recombined with genes encoding the NTDs of response regulators. Consistent with this possibility, the alignment coverage after our initial BLAST search included only 52% of the CTD sequence. Alternatively, the HTH_4_ and wH domains could share a common ancestor that is difficult to robustly infer from the isolated, divergent CTD sequences. In this case, searching with complete sequences (NTD+CTD) produced statistically significant alignments that correctly suggested an evolutionary relationship between alternatively folded CTDs. Indeed, the second phenomenon was proposed for both the Cro proteins^23–25^ and bacterial NusG transcription factors^42^.

To further discriminate whether our initial FixJ_PDB_/KdpE_PDB_ HTH_4_/wH match indicated a true evolutionary relationship or resulted from faulty homologous overextension, we next used full-length FixJ_PDB_ to query the PDB with 3 rounds of PSI-BLAST^39^, an iterative algorithm that identifies conservation patterns among homologous protein sequences. This algorithm identified stronger conservation patterns between sequences encoding HTH_4_ and wH folds and shifted the alignment registers of the CTDs. Accordingly, 97% of the FixJ_PDB_ sequence aligned with KdpE_PDB_ with an evalue of 6*10^−39^ (**Figure 1b**, right), supporting the hypothesis that the HTH_4_ and wH folds of their CTDs are distant homologs rather than alignment artifacts. Furthermore, the CTDs of 11 of the top 20 PSI-BLAST matches from this search assumed the same winged wH as KdpE_PDB_, whereas the other 9 assumed the same HTH fold as the FixJ_PDB_ query (**Table S1**). A reciprocal, three-round PSI-BLAST search using the KdpE_PDB_ sequence as query aligned 90% of this protein with FixJ_PDB_, with an e-value of 10^−29^. Notably, sequences of isolated DNA-binding domains with HTH folds were matched with the CTD of KdpE_PDB_ (wH), and sequences of isolated DNA-binding domains with wH folds were matched with the sequence of FixJ_PDB_’s CTD (HTH_4_, **Table S2**). Together, these results indicate that: (1) HTH_4_ and wH domains share a common ancestor^38^ and (2) the use of full-length sequences in our analyses, rather than isolated domains, is both legitimate and necessary to identify the relationship. Thus, all subsequent searches used full-length sequences as queries, unless otherwise noted.

Further examination of the aligned FixJ_PDB_ HTH_4_ and KdpE_PDB_ wH folds revealed regions of structural similarity and dissimilarity: both folds share a trihelical core (**Figure 1c**). By contrast, striking regions of dissimilarity are evident between (1) FixJ_PDB_’s α-helical inter-domain linker and KdpE’s corresponding quadruple-stranded β-sheet; long gaps in this alignment suggest that KdpE_PDB_’s linker region was extended through an insertion, and (2) FixJ_PDB_’s C-terminal helix aligned with KdpE_PDB_’s C-terminal β-hairpin “wing” (**Figure 1c**); the ungapped alignment of this region suggests that one of these two secondary structures may have evolved into the other through stepwise mutation.

### Alignments between response regulator sequences with HTH_4_ and wH folds indicate evolved fold switching

To further test whether stepwise mutations may have engendered a switch from α-helices to β-sheets (or vice versa), we next used an iterative Hidden Markov Model (HMM)-based alignment approach. HMM alignments are typically more sensitive than PSI-BLAST^43^ and may better avoid homologous overextension^41^. To that end, sequences for 23 non-redundant, full-length response regulators with HTH_4_ (11) and wH (12) domains were identified from the PDB. Using jackhmmer^43^, each sequence was used to query a non-redundant database of protein sequences with solved structures, including 30/46 HTH/wH domains from response regulators. These 23 full-length response regulators clustered into two subfamilies based on their CTD architectures (HTH_4_ and wH, **Figure 2a**), indicating that CTDs in the same fold families have closer evolutionary relationships than those in different fold families (**Figure S1**). Nonetheless, the C-terminal helix of the HTH_4_ domain consistently aligned with part of the C-terminal β-hairpin wing of the wH fold (*e.g*., **Figure 2b**). Furthermore, the α-helical interdomain linkers of the HTH_4_ consistently aligned with the four N-terminal β-strands of the wH domain. These results are consistent with the PSI-BLAST alignment (**Figure 2b**), suggesting that the linker may have been extended/shortened through an insertion/deletion, whereas stepwise mutation may have induced a structural interconversion between the C-terminal α-helix of the HTH_4_ and the C-terminal β-sheet of the wH.

**Figure 2.**
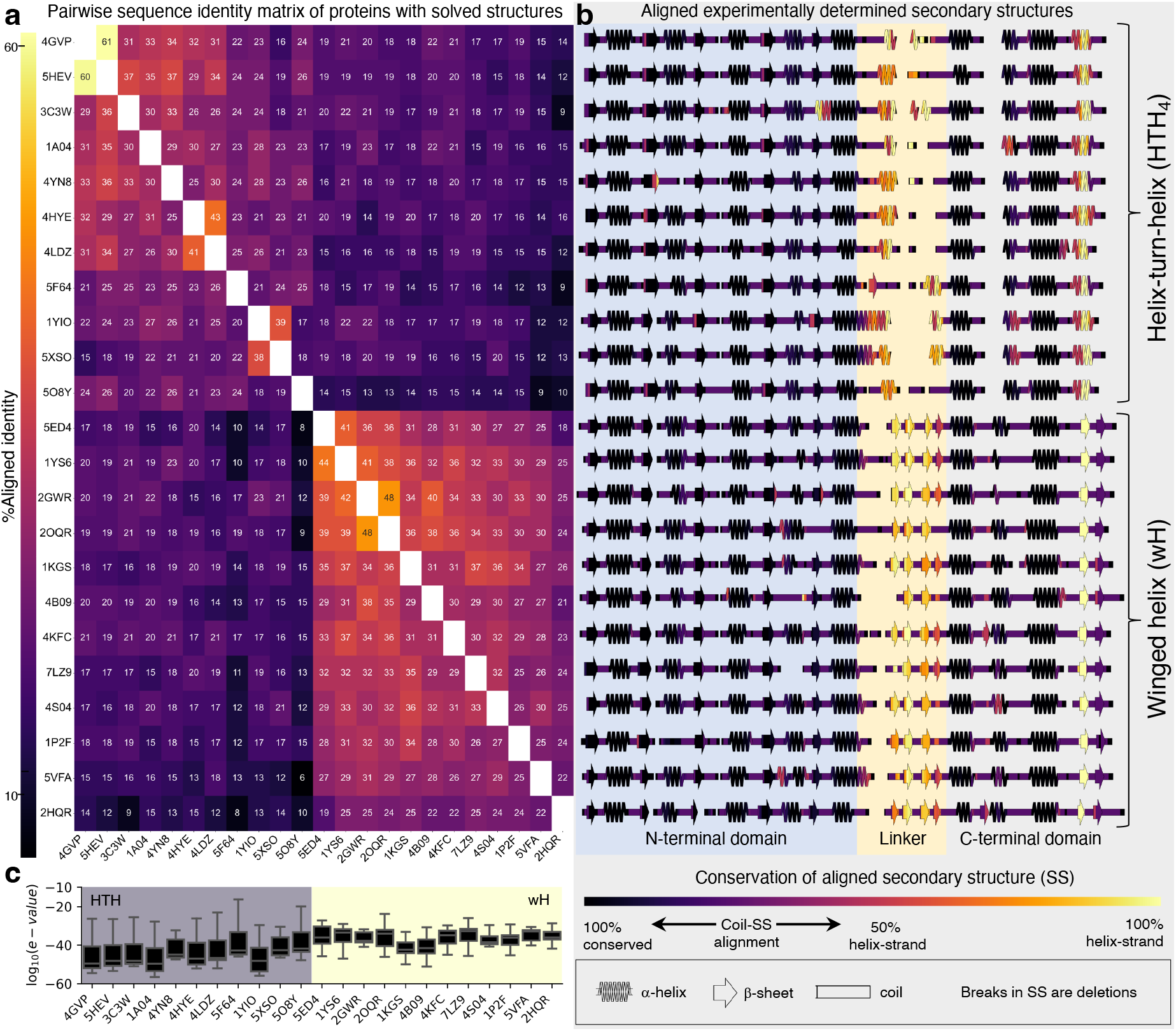
Alignments between experimentally determined HTH_4_ and wH folds consistently indicate evolved fold switching. (a). Jackhmmer-aligned sequences of response regulators with experimentally determined structures (PDB IDs) were used to calculate pairwise sequence identities. Sequences cluster into two subfamilies, with HTH_4_ (upper right bracket) and wH (lower right braket) C-terminal domains. Each row reports % aligned identities (numbers within boxes) calculated from pairwise comparisons. Identical sequences are white; all others are colored by % identity (left colorbar). (b). Experimentally determined secondary structures of each sequence in (a). The NTD, linker, and CTD are indicated by different background colors. The secondary structures are colored by their sequence-based, secondary-structure alignments with the alternatively-folded structures (HTH_4_ aligned with wH and vice versa). Identical secondary structures that consistently align are dark purple (e.g., helices that always align with helices); secondary structures that align with regions of random coil range from light purple to pink; α-helices that align with β-sheets and vice versa are colored from pink to yellow, depending on whether the alignment is more or less frequent. (c) Box and whisker plots of log_10_(e-values) of jackhmmer searches of sequences that used one fold to query sequences from the alternative subfamily (HTH_4_ against wH or vice versa). The distributions of each HTH_4_ (gray background)/wH (yellow background) box were derived from n ∈ [7,12] e-values; each box bounds the interquartile range (IQR) of the data (first quartile, Q1 through third quartile, Q3); medians of each distribution are gray lines within each black box; lower whisker is the lowest datum above Q1-1.5*IQR; upper whisker is the highest datum below Q3 + 1.5*IQR.

The possible relationship between HTH_4_ and wH folds was further supported by assessing the e-value distributions from alignments between the full-length proteins with (1) homologs from their own subfamily and (2) homologs from the alternatively-folded subfamily (**Figure 2c**, gray/yellow backgrounds, respectively). Median e-values of the alignments between the sequence of a given experimentally determined fold (HTH/wH) and the set of sequences with the alternative fold (wH/HTH) ranged from e-33 to e-43, suggesting significant evolutionary relationships across all members of the two subfamilies (**Figure 2c**). As expected, the median e-values among sequences of similar folds ranged from e-54 to e-72 (**Figure S2a**), indicating closer evolutionary relationships.

Statistically significant alignments were also identified between full-length query sequences and isolated CTDs with the alternative fold in 22/23 response regulators. Median e-values of these alignments ranged from e-04 to e-09, whereas median e-values of aligned sequences from the same fold family ranged from e-17 to e-30 (**Figure S2b**). These domain-specific alignments further support the evolutionary relationship between HTH_4_ and wH domains.

### Phylogenetic analyses of HTH_4_ and wH proteins

Although these structure-based sequence searches were consistent with evolved fold switching in the C-terminal region of HTH_4_ and wH domains, the mechanism of secondary structure conversion was obscured by the alternative locations of sequences inserted into the CTD of the longer wH homologs. PSI-BLAST fully aligned the C-terminal α-helix of the HTH_4_ with the β-hairpin of the wH (**Figure 1b**), suggesting a full secondary structure conversion. By contrast, jackhmmer aligned the C-terminal α-helix of the HTH_4_ with only the first β-strand of the wH (**Figure 2b**), suggesting a partial conversion along with an insertion. To discriminate between these options and to identify possible evolutionary trajectories between the fold families, we next collected a large set of response regulator sequences with HTH_4_ and wH output domains. To that end, the FixJ_PDB_ and KdpE_PDB_ sequences were queried against the nr database using protein BLAST to identify 581,791 putative homologs. Given the size of this sequence set, we developed several strategies for curating and sampling the data (**Methods**) so that the final subset of sequences would be small enough for various phylogenetic analyses but large enough to adequately represent the large family of response regulators.

To that end, the 581,791 sequences were grouped into 367 clusters using a greedy clustering algorithm and filtered to 85% redundancy for a final number of 23,791 sequences. Clusters were then compared to identify 13,006 FixJ-like sequences and 10,785 KdpE-like sequences. Sequences within each group readily aligned; however, the two groups had overall low sequence identities with each other. Several approaches were attempted to align these groups. One attempt identified a “transitive homology pathway” of 7 sequences connecting HTH_4_ to wH sequences (**Table S3**) that was used to match the FixJ-like (HTH_4_) and KdpE-like (wH) alignments. However, when a phylogenetic tree was constructed in IQ-Tree for the combined set of 23,791 sequences, its quality was poor (*i.e*., 140 gaps/360 positions in the KdpE_PDB_ sequence) and failed to converge after 3 rounds of bootstrapping.

Nevertheless, the transitive homology path suggested the existence of additional sequences that might bridge the HTH_4_ and wH fold families. Thus, we searched the original sequence set with an alternative approach. First, we categorized clusters with ≥100 sequences by their CTD architectures to identify 74,741/387,276 sequences with HTH_4_/wH output domains. These sequence sets were used to construct BLAST libraries. Next, the sequences with HTH_4_ output domains were filtered to 50% redundancy, and the remaining 4,520 sequences were queried against the wH library with protein BLAST. If a match was statistically significant, we searched NCBI sequence records for CTD structure annotations of both sequences, which are typically inferred from Hidden Markov Models. These results were used to distinguish BLAST matches between different fold families (sequence pairs with 1 HTH_4_ and 1 wH) from matches from the same fold family. Sequence pairs with annotations from different fold families were retained; this process identified 3136 matches between 664 HTH_4_ and 2541 wH proteins with mean/median e-values of 4*10^−10^/5*10^−16^. Reciprocal BLAST searches, using the wH sequences as queries, were successfully performed in all 3136 cases, with mean/median e-values of 1*10^−8^/2*10^−16^; these higher e-values likely reflect the smaller size of the HTH_4_ database or the longer lengths of wH sequences relative to HTH_4_.

Next, we aligned the 3,205 sequences using two different methods, Clustal Omega^44^ and MUSCLE^45^ (**Supplementary Data 1**). Again, a key difference between these cross-family multiple sequence alignments (MSAs) was the location of sequences inserted into/deleted from the longer wH/shorter HTH_4_ homologs. Nevertheless, in both cross-family MSAs, the C-terminal helix of the HTH_4_ aligned fully with the C-terminal β-sheet wing of the wH, indicating evolution from α-helix to β-sheet by stepwise mutation rather than insertion or deletion (**Figure 3a, Figure S3**). In the Clustal Omega alignment, a two-residue gap found in >99% of HTH_4_ folds was also found in an annotated wH fold (wH_wing_gap_), further suggesting that the α-helix ↔ β-sheet interconversion occurred through stepwise mutation. Furthermore, several HTH_4_ sequences with linker lengths similar to wH sequences were identified (*e.g*., HTH_4_insert_ in **Figure 3a**), demonstrating that long linkers are not exclusive to wH folds. Sequences within the alignment were diverse, with mean pairwise identities of 31% among HTH_4_ folds, 40%among wH folds, and 31% across folds. Notably, evolutionary conservation patterns differed between the HTH_4_ and wH folds (**Figure S4**). Particularly, the C-terminal helix of the HTH_4_ did not show strong conservation patterns, whereas the β-strand wing of the wH did. As suggested by Cordes and colleagues^24^, such distinct conservation patterns may explain why homology between isolated wH and HTH_4_ sequences was not detected in the PSI-BLAST and jackhmmer searches against the PDB.

**Figure 3.**
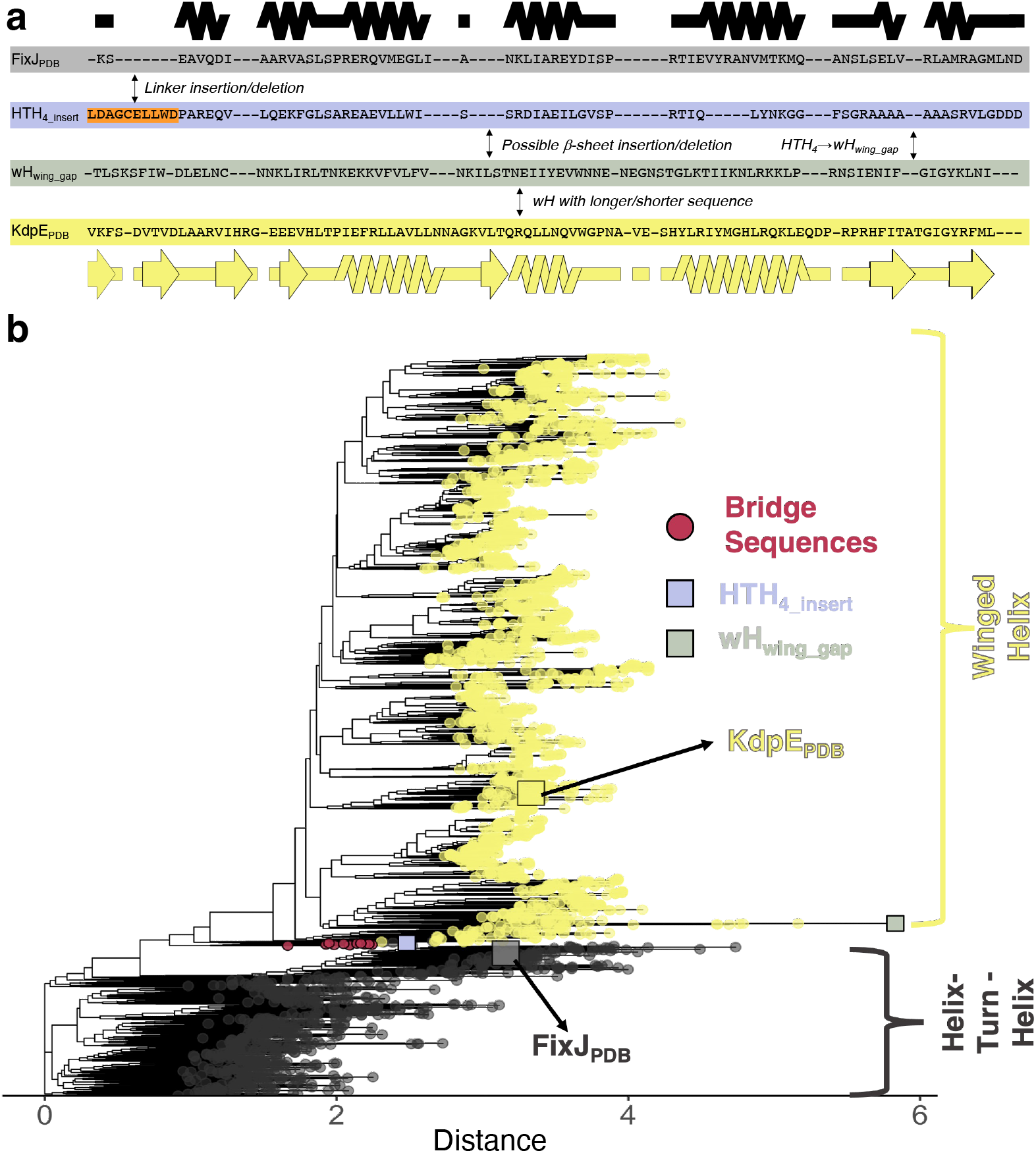
(a) Clustal Omega alignment of 3,205 HTH_4_ and wH sequences indicates complete conversion of C-terminal secondary structure over evolutionary history. Secondary structure diagrams were generated using the structures of FixJ_PDB_ (black) and KdpE_PDB_ (yellow). Background colors of the four sequences match those in the phylogenetic tree. Notes in the spaces between sequences show important changes: (1) orange linker insertion (or deletion, depending upon the properties of ancestral sequences) (2) fold conversion (3) sequence elongation/deletion. The word in front of a slash represents what happens if a sequence changes from top to bottom; the word following the slash represents what happens if a sequence changes from bottom to top. A common ancestor between the FixJ_PDB_ and KdpE_PDB_ sequences is also possible. b) Maximum-likelihood phylogenetic trees suggest an evolutionary path between response regulators with HTH_4_ and wH folds. Sequences with CTDs annotated as HTH/wH from NCBI protein records are gray/yellow. The clade containing the 12 identified bridging sequences is highlighted in pink. HTH_4_insert_ provides an example of an annotated HTH_4_ sequence whose linker length was similar to wH; wH_wing_gap_ provides an example of a wH sequence with a 2-residue deletion similar to those found in >99% of the C-terminal helices of aligned HTH_4_ sequences. Distance units are arbitrary, though sequences further in space have more distant evolutionary relationships.

Finally, we generated a bootstrap-supported, phylogenetic tree for the cross-family MSA. Strikingly, results revealed a sequence clade that appears to bridge the two fold families (**Figure 3b, Figures S5**). The 12 sequences of this clade include one identified in the transitive homology path; all 12 have output domains annotated as HTH_4_ and originated from several bacterial phyla (**Table S4**). In the phylogenetic tree, these 12 sequences adjoin branches with wH and HTH_4_ CTDs (**Figure 3b**), suggesting that their ancestors might be evolutionary intermediates between the two folds. To assess the statistical robustness of the HTH-bridge-wH interface, we quantified the frequency of its occurrence using trees rooted in all 6393 possible branch points. The log-likelihood of each rooted tree was calculated using the approximately unbiased test^46^ (p-AU, **Figure S6A**). Of the 6393 possible rootings, 18 had a p-AU score ≥ 0.8 (**Figure S6B**), indicating statistical significance. In all 18 cases, the bridge sequences adjoined branches with annotated wH and HTH_4_ domains (**Figure S7**), strongly supporting the role of this clade as an evolutionary bridge between the two folds.

### A mutational pathway between two folds

We next examined the predicted structural properties of sequences in the bridge clade. To that end, structural models of each bridge sequence were produced with AlphaFold2^14^ (AF2). Strikingly, all models assumed the HTH_4_ fold (**Figure S8**). This result suggests a few possibilities. First, some bridge sequence(s) might interconvert between HTH_4_ and wH folds; previous work has shown that AF2 generally predicts only one dominant conformation of proteins that can switch between two folds^47,48^. Second, the AF2 predictions could be unreliable, and some or all bridge sequences could, in fact, assume wH folds. Thirdly, the fold transition might have occurred in earlier ancestors located at nodes linking most HTH_4_ and wH sequences. These nodes connect the two fold families in the tree (**Figure S5**), suggesting that their corresponding ancestral sequences may have had properties of both HTH and wH folds.

Thus, we next performed ancestral sequence reconstruction and generated additional AF2 models for the ancestral sequences bridging the HTH_4_ and wH folds (**Figures 4, S5)**. Note that the linkers of all ancestral sequences were as long as the wH linkers. Our rationale was that the linkers of some HTH_4_ sequences near the bridge region were equally long as the linkers of wH sequences (**Figure 3, Figure S3**), suggesting that these linkers may have already been modified by a large insertion.

**Figure 4.**
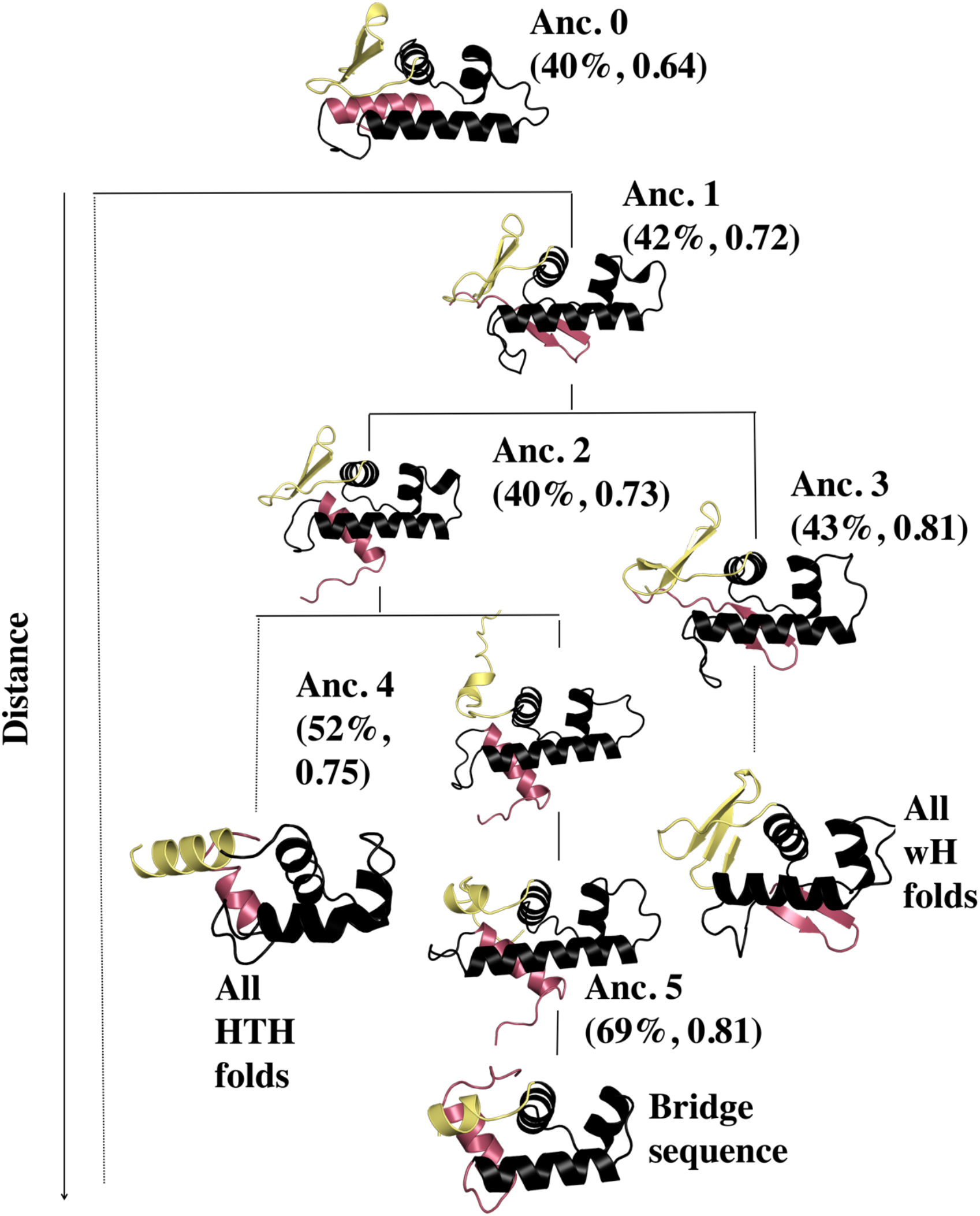
AlphaFold2 predictions for the CTDs of the reconstructed ancestors appear to switch folds in response to 2-3 mutations in the most C-terminal secondary structure element. The earliest ancestor appears to be the longer version of an HTH_4_, from which wH folds evolved. The fold-switching C-terminal helix/β-hairpin is shown in pink, and the structurally plastic linker is shown in yellow. The bridge sequence used in this plot was TME68356.1, the one nearest the ancestral node in **Figure 3b**.

Intriguingly, results from ancestral reconstruction suggest that the ancestor sequences may have had structurally plastic regions that could switch between α-helices and β-sheets in response to mutation (**Figure 4, Table S5**). Notably, Ancestor 0’s most C-terminal secondary structure element is an α-helix, Ancestor 1’s is a β-hairpin, and Ancestor 2’s switches back to an α-helix (**Figure 4**, pink). Interestingly, the sequence of Ancestor 1’s β-hairpin is 83% identical to the sequences of both Ancestor 0’s and Ancestor 2’s C-terminal helices, which are 75% identical to one another. These results suggest that just two mutations can switch the C-terminal α-helix to a β-sheet and back again through a different set of sequence substitutions.

The N-terminal linker region (**Figure 4**, yellow) also appears to be plastic. In Ancestors 0-2, this linker is partially folded into a β-hairpin structure, whereas in Ancestor 3 the linker assumes a fully folded 4-β-sheet structure. By contrast, the linker assumes a partially helical structure in Ancestors 4-5 and in the modern-day bridge sequence (**Figure 4**).

Taken together, these results suggest that ancestors of sequences in the bridge clade may have had propensities for *both* wH and HTH_4_ folds. To further test this possibility, both PSI-BLAST and hmmer searches were carried out between the ancestral CTD sequences and PDB structures with both HTH_4_ and wH folds. Statistically significant cross-fold matches were identified in all cases except for Anc. 3 (**Supplementary Data 2**). By comparison, the earlier PSI-BLAST and hmmer searches of the isolated CTDs of existing HTH_4_ and wH sequences matched homologs with the same but not the alternative fold.

### Evolution from HTH_4_ to wH may have expanded DNA binding specificity

Finally, we sought to identify whether the shift from HTH_4_ to wH folds may have had some evolutionary advantage. Examination of experimentally determined HTH_4_ and wH response regulator structures in complex with their cognate DNA partners suggests that one benefit of the structural transformation might have been expanded binding specificity. On average, the HTH_4_ folds contact 17 unique nucleotides, whereas the wH folds contact 22 (**Figure 5a**). Both HTH_4_ and wH folds have a single recognition helix that binds the major grove, and the C-terminal β-hairpin of winged helices also contacts the minor groove (**Figure 5b**). As such, wH domains can likely recognize more unique nucleotide sequences than HTH.

**Figure 5.**
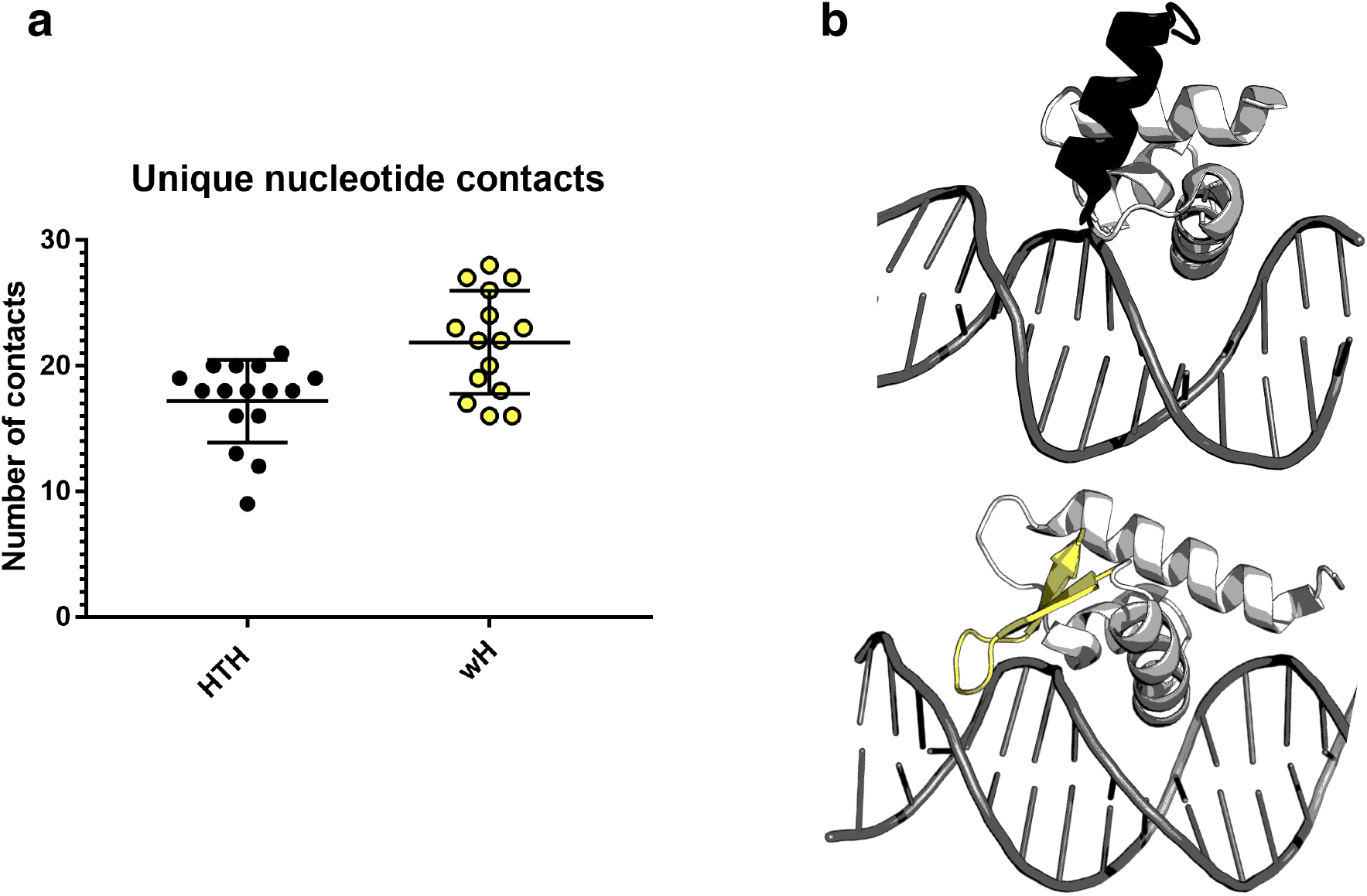
HTH_4_ domains contact fewer nucleotides, on average, than wH domains. (a) Simplified box-and-whisker plot with overlaying datapoints for the number of contacts between HTH_4_ and DNA (black) and wH and DNA (yellow). On average, HTH_4_ domains have 5 fewer DNA contacts than wH domains. (b). Examples of DNA (gray) interactions with HTH_4_ and wH domains, above and below, respectively. The C-terminal α-helix of the HTH_4_ (black, above) does not contact the DNA, whereas the β-hairpin wing of the wH (yellow, below) contacts the minor groove. Structurally similar parts of the HTH_4_ (PDB ID: 1h0m, chain D) and wH (PDB ID: 4hf1, chain A) folds are light gray. This result and the corresponding increase in the possible number of unique DNA sequences that could be recognized by the wH might explain why it evolved from the HTH_4_ in response regulators.

### Conclusions

Decades of research suggest that protein secondary structure is largely conserved over evolutionary history^49,50^. Here, we show how evolutionary sequence variation has morphed an α-helix into a β-sheet in bacterial response regulators. A variety of studies have shown previously that new protein folds can evolve through various mechanisms, such as insertions, deletions, and circular permutation^51^. Others have shown that (1) proteins with conserved secondary structures can evolve different tertiary arrangements^21,22,52^, (2) a basis set of homologous protein fragments, known as words, is conserved in disparate protein structures^53^, and (3) segments of structure (“themes”) can be mixed and matched to form different protein architectures^54–56^.

In addition, previous work reported that sequence substitutions can switch secondary structure of existing proteins and hypothesized that similar secondary structure changes may have occurred over evolutionary history^19,25,57,58^. Our work confirms these hypotheses by identifying a statistically significant evolutionary trajectory between two protein folds in the response regulator CTDs that switch from α-helix to β-sheet. These findings are supported by ancestral sequence reconstruction, structural models, and several sequence alignment methods. Furthermore, this evolved fold switching likely had a functional consequence: expanding DNA-binding specificity. Notably, HTH_4_ and wH folds are not limited to the superfamily of response regulators. In other families, the wHs could have evolved from HTH or HTH_4_ ancestors through similar or additional mechanisms (and the evolutionary order may differ).

Structural transformations, such as the one identified here, may be more common in the evolutionary record than currently realized. Detecting such transformations has been impeded by: 1) limited abundance and diversity of sequenced genes, (2) limited availability of experimentally determined protein structures, and (3) assumptions underlying the statistical methods used to infer homology. Here, we overcame these barriers by searching a deep and diverse set of homologous sequences, which were unavailable decades ago when related studies were performed^25,38,59^. Our results also highlight the challenge in finding putative evolutionary pathways between two folds. First, we established the necessity of using full-length–rather than domain-limited–sequences to compare response regulators with HTH_4_ and wH domains. Then, within the ~600,000 sequences of the large superfamily, we identified the 0.5% of protein sequences with high sequence identities but distinct folds. Among these, an evolutionary pathway from HTH_4_ to wH was consistently observed, with a clade of “bridge sequences” occupying a key location in the pathway. Notably, these bridge sequences were identified from metagenomic sequencing performed primarily in 2018 and 2019, which demonstrates the importance of new sequencing techniques and initiatives for advancing evolutionary studies^60^ and suggests that more instances of evolved fold switching might now be identifiable.

## Methods

### BLAST and PSI-BLAST searches of the PDB

To identify the putative evolutionary relationship between FixJ_PDB_ and KdpE_PDB_, we performed protein BLAST searches with maximum e-value of 1e-04 on all sequences within the Protein Data Bank (PDB) against all other PDB sequences, as previously reported^16,40^. To determine whether homologous sequences folded into different structures, secondary structure annotations of each PDB, by DSSP^61^, were aligned in register with their corresponding BLAST alignments and compared one-by-one, position-by-position. This approach allowed us to quantitatively assess the similarity of aligned secondary structures. A potential match was required to have a continuous region of at least 15 residues in which at least 50% of the residues showed α-helix ↔ β-sheet differences. Using this approach, the sequence of FixJ_PDB_ matched the sequence of KdpE_PDB_ with an e-value of 1e-07; differing secondary structures in the C-terminal output domains were identified through DSSP comparison. Subsequent three-round PSI-BLAST searches of FixJ_PDB_ and KdpE_PDB_ sequences against all PDB sequences were performed with a gap open penalty of 10 and a gap extension penalty of 1. In CTD PSI-BLAST searches, the sequences for FixJ_PDB_ and KdpE_PDB_ spanned residues 124-205 and residues 129-225, respectively.

### jackhmmer alignments of structures with response regulator sequences

To test the PSI-BLAST results obtained previously, jackhmmer searches were also performed on HTH_4_ and wH sequences with experimentally determined structures. Accordingly, structures of 23 full-length response regulators with HTH_4_ (11) and wH (12) output domains were identified from the Evolutionary Classification of Protein Domains (ECOD) database^62^. Five rounds of jackhmmer were run on each of the 23 sequences with gap open/extension probabilities of 0.05 and 0.5, respectively, using a database of all sequences downloaded from the PDB (7/15/2021) with sequence duplicates removed. Sequence identities from each row of **Figure 2a** were calculated from each sequence alignment generated by jackhmmer run on the sequence of the PDB entry with ID labeling each respective row.

DSSP annotations were aligned in register with each jackhmmer-generated sequence alignment to compose the secondary structure diagrams in **Figure 2b**. In further detail, secondary structure annotations of each of the 11 HTH_4_s were compared with secondary structure annotations of 48 wHs identified from ECOD; likewise, secondary structure annotations of each of the 12 wHs were compared with secondary structure annotations of 35 HTH_4_s identified from ECOD (**Supplementary Data 3**). Similarities of each pair of aligned secondary structure (46 pairs for each of the 11 HTH_4_ proteins, 30 pairs for each of the 12 wH proteins) were scored as follows: +1 for a position with identical secondary structures (helix:helix [H,G,I in DSSP notation] or strand:strand [E in DSSP notation]) and −1 for a position with alternative secondary structures (helix:strand or strand:helix using the same DSSP notations as above). Position-specific scores were normalized by the frequency of ungapped residue pairs in each position, including coil-secondary structure alignments, effectively scored as 0. These normalized position-specific scores were used to generate the colormaps of each secondary structure diagram.

### Identifying large sets of response regulators’ genomic sequences

The full sequences of both FixJ_PDB_ (PDB ID 5XSO, chain A) and KdpE_PDB_ (PDB ID 4KFC, chain A) were searched against the nr database (10/8/2020) using protein BLAST with a maximum e-value of 1e-04 and a maximum of 500,000 alignments per search. Full sequences from each alignment were retrieved by their NCBI accession codes using blastdbcmd on the nr database. All sequences from both searches were combined, which totaled 999,912 after sequence duplicates were removed. Sequences with either fewer than 162 or more than 300 residues were removed because they likely lacked the proper response regulator domain structure, leaving 581,791 sequences. This was too many to curate using standard tools, and many sequence identities were well below the ~40% identity threshold, below which many alignment tools become unreliable^63^. Thus, to further analyze these sequences, we performed the clustering and sampling methods described in the following sections.

### Generating sequence clusters

From set of 581,791 sequences, a basis set of 367 sequences – each with <24% pairwise identity to all other members of the set – was selected to seed sequence clustering. Above this threshold, response regulator sequences would be expected to assume similar structures^49^. To identify this set of seed sequences, the first sequence in the list of 581,791 sequences (FixJ_PDB_) was chosen. Subsequent sequences were aligned with FixJ_PDB_’s sequence using Biopython^64^ pairwise2.align.localxs with gap open/extension penalties of −1, −0.5, respectively. If a sequence’s pairwise identity with the FixJ_PDB_ sequence <24%, it was added to the basis set. Sequences in the list were aligned with all sequences previously added to the basis set and included only if the identities of *all* pairwise alignments were <24%, yielding 367 total basis sequences. The remaining 581,424 sequences were clustered with the basis sequence to which they had the highest aligned pairwise identity, determined exhaustively by aligning all sequences with all basis sequences using pairwise2.align.localxs, with parameters as before.

To further reduce the total number of sequences, we disregarded the 251 clusters with fewer than 50 sequences. The remaining 116 clusters comprised 103 “medium” clusters (<5000 sequences) and 13 “large” clusters (>4000 sequences). Of the large clusters, one contained the sequence of FixJ (PDB ID 5XSO) and 283,762 other sequences, and another contained the sequence of KdpE (PDB ID 4KFC) and 25,035 other sequences.

### Curating sequence clusters

#### Medium clusters

Sequences within each medium cluster were first aligned using Clustal Omega^44^. Visual inspection revealed that some alignments were biased by sequences that were either substantially shorter or longer than majority of the homologs in their cluster. To computationally identify and filter out such sequences, we identified (i) “sparse zones” by searching for windows of 8 positions where more than 95% of the sequences contained gaps, and (ii) “populated zones” by searching windows of 10 positions where more than 90% of the sequences contained amino acid residues. Sequences with (1) ≥ 10% of their amino acids in sparse zones or (2) < 10% of their amino acids in populated zones were removed from the cluster. The 10% thresholds were determined empirically to best perform this “culling” step. Next, we performed ~2-7 successive iterations of culling and Clustal Omega alignments, until the number of sequences in each cluster converged. During this process, 9 medium clusters shrunk to fewer than 50 sequences and were subsequently ignored, leaving 94 medium clusters.

Finally, since Clustal Omega’s global alignment algorithm does not accurately report phylogeny or suggest structure, the multiple sequence alignments were further aligned using PROMALS^65^, which first groups sequences based on phylogeny and then performs local alignment of recognized structural domains. The quality of all cluster alignments was inspected visually.

#### Large clusters

The large clusters, with thousands of sequences, required different strategies to appropriately generate a subsample that was tractable for additional sequence analyses. To determine subsample sizes that adequately represented the sequence composition within clusters, three independent, random subsamples of 1000 and 5000 sequences were extracted from the FixJ cluster, and three 5000 sequence subsamples were extracted from the KdpE cluster. These subsamples were subjected to iterative culling and alignments like the medium clusters (described above).

Next, the multiple sequence alignments (MSAs) of these subsamples were uploaded to ConSurf^66^ (https://consurf.tau.ac.il/consurf_index.php). Resulting scores were compared to determine how many sequences were required to give consistent evolutionary rates. Results indicated that 5000 sequences were required for an adequate representation of the both the FixJ and KdpE clusters. Visual inspection of heatmaps generated from sequence identity matrices of these sequence alignments supported the conclusion that 5000 sequences evenly sampled the sequence space. Thus, to represent the FixJ and KdpE clusters, we randomly chose one of its 5000 subsamples sequence sets. For 8 of the 11 large clusters with >5000 sequences, we similarly subsampled 5000 sequences. The 3 large clusters with <5000 sequences were curated as described for the medium clusters.

### Constructing FixJ and KdpE-specific MSAs

The high sequence diversity between clusters, with cross-cluster pairwise aligned sequence identities often <24%, impeded MSA assembly of the FixJ-KdpE superfamily. Thus, we looked for strategies to assemble sequences from the 94 medium clusters, 11 large cluster subsamples, and the 5000-sequence subsamples of the FixJ and KdpE large clusters into one combined MSA. First, we classified the clusters into two half-families with sequences resembling those in either the FixJ or KdpE large clusters. To that end, we matched sequences from each cluster with all sequences from the FixJ and KdpE large clusters with protein BLAST. Sequences from these clusters tended to align with high statistical significance to one of the large clusters but not both, simplifying cluster classification. This approach showed promise because sequences from each cluster aligned to sequences from other clusters with identities ≥38%, fostering reliable alignments. After completing all BLAST searches, 45 medium and 6 large clusters were assigned to the FixJ half-familiy for a total of 13,006 sequences and 49 medium and 5 large clusters to the KdpE half-family for a total of 10,785 sequences.

Despite sampling and curation, both half-families were too large to create an MSA using conventional tools. Thus, we used an alternative approach in which two reference alignments were generated using Clustal Omega to align representative sequences from each cluster (51 sequences for FixJ and 54 for KdpE). PROMALS was then used to refine the two half-family reference MSAs. Upon visual inspection, 7 sequences were removed from the KdpE reference MSA because they generated many gaps in the alignment; their clusters of origin were subsequently ignored. The remaining sequences in the KdpE reference MSA were realigned using Clustal Omega and PROMALS. Finally, upon visual inspection, the registers of prolines and charged amino acids were manually edited to match in 3 sequences (PSQ94266, HBD38673, KEZ75144) between the registers 225 and 270 in the KdpE reference MSA. No such manual curation was needed in the FixJ MSA. Sequences within each of the remaining 98 clusters were then (i) independently aligned with PROMALS and (ii) integrated into the appropriate half-family reference MSA using MARS (Maintainer of Alignments using Reference Sequences for Proteins^67^). The MARS program allows curated sequence alignments with at least one sequence in common to be merged with each other without re-aligning the whole sequence set. Using this program, all sequences of the 51 FixJ-matching clusters and the curated subsample of the FixJ cluster were merged, using the FixJ half-family reference MSA as a guide. Similarly, all sequences of the 47 KdpE-matching clusters along with the curated subsample of the KdpE cluster were merged.

### Constructing a FixJ-KdpE superfamily MSA

The pairwise identities of sequences across the two half-families were too low to reliably create an MSA. Thus, we tried a “transitive homology” approach to combine the half-family alignments into one alignment for the superfamily. First, we identified a “path” of related sequences^68,69^ following the logic that, if sequences A and B are homologous and sequences B and C are homologous, then homology between sequences A and C can be assumed through the “bridge” sequence B. To carry out this strategy, we used protein BLAST to search for the highest sequence identity match between the unsampled FixJ and the KdpE large clusters (i.e., the clusters with >250,000 and >25,000 sequences). This hit was then queried against the database of the opposite fold and so on until we identified 7 sequences with pairwise sequence alignments ≥38% that connected the FixJ sequence to the KdpE sequence (**Table S3**). Note that the “bridge” sequence TME68356 (**Table S4**) could align well with another sequence in either half-family, although it was originally assigned to the KdpE half-family. The top/bottom four sequences in **Table S3** were aligned with the FixJ/KdpE half-families using Clustal Omega. We next used MARS to combine half-family alignments using the bridge sequence as the reference. The resulting whole family MSA contained 45,199 sequences. These sequences were filtered to 85% redundancy with CD-HIT, ultimately yielding an MSA with 23,791 sequences. However, when a phylogenetic tree was constructed in IQ-Tree for this sequence set, its quality was poor (*i.e*., 140 gaps/360 positions in the KdpE_PDB_ sequence) and failed to converge after 3 rounds of 1000 bootstrapping iterations each.

### Constructing a cross-family MSA

The transitive homology path identified above (**Table S3**) suggested the existence of additional sequences that might bridge the HTH_4_ and wH folds. Accordingly, the five/six previously-assigned FixJ/KdpE sequence clusters with >4000 sequences were each combined and converted into two BLAST databases representing HTH_4_ (FixJ-like) and wH4 (KdpE-like) sequences. Sequences within the combined FixJ sequence clusters were reduced to 50% redundancy using CD-HIT^70^ with a word size of 2, as recommended. Protein BLAST searches were performed on each of the remaining 4,520 sequences with a maximum e-value of 1e-04 using the full KdpE_PDB_ database. All 8607 alignments with minimum sequence identities and lengths of 33% and 200 residues, respectively, were considered significant. To ensure that these alignments truly matched HTH_4_ with wH sequences, NCBI records of 1,793 HTH_4_ and 4,995 wH sequences were retrieved using NCBI’s efetch. Each record was searched for structural annotations of its CTD (HTH or wH). Ultimately, 3,074 BLAST matches, each with one annotated HTH and one annotated wH CTD were retained.

To identify additional HTH sequences that might match with wH sequences, additional BLAST searches were run on all 4 HTH_4_ sequences in our set of 3,074 matches that aligned with wH sequences with ≥38% pairwise identity. This time, the database comprised all 581,791 length-limited sequences identified from the initial FixJ and KdpE BLAST searches. These searches, intended to identify additional HTH_4_ sequences regardless of how they were clustered, yielded 66 putative HTH sequences that might match well with additional wH sequences. Finally, 66 additional Protein BLAST searches were performed by querying each of the 66 putative HTH sequences against all sequences from the 47 KdpE-matching clusters identified previously. The resulting 62 matches with minimum sequence identities and lengths of 33% and 200 residues and HTH/wH annotations from their NCBI records, identified as before, were included, totaling 3,136 matches between 3,203 sequences. For reference, the sequences of FixJ_PDB_ and KdpE_PDB_ were also included; these two sequences had minimum aligned identities and lengths of 32% and 198, respectively, to sequences encoding the alternative folds.

The resulting 3,205 sequences were aligned in two ways, with Clustal Omega and with MUSCLE^45^ version 3 using the super5 command. Columns with >75% gaps were removed from both alignments for further analyses. The final alignments showed full overlap between the C-terminal helix of the HTH_4_ and the β-hairpin wing of the wH. Subsequent phylogenetic analyses and ancestral sequence reconstruction were performed on the Clustal Omega alignment.

### Conservation scores and rate of evolution

A version of ConSurf that could be run locally, Rate4Site^71^ (https://www.tau.ac.il/~itaymay/cp/rate4site.html), was used to compute evolutionary rates for the full alignment of 3,205 sequences as well as the separate HTH_4_ and wH subfamilies (664 and 2541 sequences, respectively; **Figure S4**). This program requires an MSA file to compute a phylogenetic tree. We chose the empirical Bayesian method to generate the rates, which significantly improves the accuracy of conservation scores estimations over the Maximum Likelihood method^71^. The scores are represented as grades ranging from conserved (9) to variable (1).

### Phylogenetic analyses of the cross-family MSA

#### Constructing a maximum-likelihood tree and performing bootstrapping

A maximum-likelihood (ML) phylogenetic tree was inferred from the alignment with FastTree^72,73^, using the Jones-Taylor-Thorton/JTT^74^ models of amino acid evolution and the CAT^75^ approximation to account for the varying rates of evolution across sites. This tree was further supported by ultrafast bootstrapping (UFBoot^76^) as implemented in IQ-Tree2^77^. We used ModelFinder^78^ to identify the best fitted evolutionary model for the MSA (chosen model - LG+F+R10), and then evaluated branch support with 1000 UFBoot replicates. The minimum correlation coefficient for the convergence criterion was set at 0.99. A consensus tree was also generated (**Figure S5**).

#### Rooting the phylogenetic tree

The ML and consensus trees generated by FastTree and IQ-Tree2, respectively, lacked information on root placement of the estimated phylogeny. Ideally, external information – such as an outgroup – is used to root the tree. However, we could not use an outgroup because it was not possible to identify a single sequence outside of our alignment that was homologous to both folds. Therefore, we combined the nonreversible model with a maximum likelihood model^79^ used to calculated the loglikelihoods of the trees being rooted on every branch of the tree. Bootstrapping of 10,000 replicates was performed to obtain reliable results. The method returns a list of 6393 trees rooted at each node and sorted by log-likelihoods in descending order, along with other scores by different tests, as follows; bp-RELL: bootstrap proportion using RELL method^80^, p-KH: p-value of one-sided Kishino-Hasegawa test^81^, p-SH: p-value of Shimodaira-Hasegawa test^82^, c-ELW: Expected Likelihood Weight^83^ and the p-AU: p-value of approximately unbiased (AU) test^46^.

The AU test uses a newly devised multiscale bootstrapping technique developed to reduce test bias and to obtain a reliable set of statistically significant trees. The AU test, like the SH test, adjusts the selection bias overlooked in the standard use of the bootstrap probability and KH tests. It also eliminates bias that can arise from the SH test^46^. Overall, the AU test has been shown to be less biased than other methods in typical cases of tree selection and is recommended for general selection problems^46^. Hence, we relied on p-AU (p-values from AU) to get a list of 18 most-likely rooted trees with p-AU > 0.8.

#### Ancestral sequence reconstruction

Ancestral sequence reconstruction was performed using maximum likelihood methods implemented in IQ-Tree2, which uses the algorithm described in Yang et al.^84^ Ancestral sequences were determined for all nodes of the consensus tree (**Figure S5**) using the empirical Bayesian method. Posterior probabilities are reported for each state (amino acid) at each node. We scored the nodes in three steps. First, we calculated the average probability considering all assigned states at the node. Then, replacing the states by the amino acids in the bridge sequence (TME68356.1), we calculated the total p-value. Finally, calculated the pairwise sequence identity between ancestral sequence and the bridge sequence. Using all three criteria, we identified 6 reconstructed sequences with low p-values near the bridge sequences. These sequences were used for downstream analysis and model building.

### Predicting structures of ancestral and bridge sequences

The FASTA sequences of the 6 reconstructed ancestors, along with the 12 bridge sequences, were used as input to the AlphaFold2.1^14^ structure prediction model. Structures ranked 0 were depicted in **Figures 4** and **S8**. To test the plausibility of the AF2-generated structures for the reconstructed ancestors and bridge sequences, we examined recently released AF2 predictions for 338 HTH_4_ and 937 wH sequences^85^. AF2 predictions matched genomic annotations in every case. Prediction qualities varied: of 1275 predicted structures, 29% were predicted with high confidence, 58% had moderate confidence, and the remaining 13% had low confidence.

### Counting protein-DNA contacts

The unique nucleotide contacts between the response regulators and their corresponding DNA sequences were identified using Resmap^86^, a tool that uses the atomic coordinates from PDB files to calculate intra-atomic distances for non-covalent interactions under set thresholds. The default distance thresholds for different interaction types that were used are: (1) Hydrogen bonds - <= 3.5 Å, (2) Hydrophobic interactions - <=4.5 Å, (3) Aromatic interactions - <= 4.5 Å, (4) Destabilizing contacts - <= 3.5 Å, (5) Ion pairs - <= 5.0 Å, (6) Other contacts (which include van der Waals interactions) - <= 3.5 Å. Since the nomenclature for DNA atoms has changed since the development of Resmap, the PDB files were manually edited to match Resmap’s input format with the following changes: (1) Symbol replacements of ‘ to *, (2) the nucleotide atoms (A,C,G, or T) were appended with the prefix ‘D’ (DA, DC, DG, DT), (3) the edited nucleotide atoms were also assigned unique atom identification numbers. The PDB files with these changes were then inputted into Resmap to identify unique contacts between of atoms in the protein chains with the atoms in DNA chains.

### Scripts and figures

Protein figures were generated in PyMOL^87^ (https://pymol.org/2/), plots and heatmap in Matplotlib^88^ (https://matplotlib.org/stable/index.html) and seaborn^89^ (https://seaborn.pydata.org/). Phylogenetic trees were visualized with ggtree (https://guangchuangyu.github.io/ggtree-book/chapter-ggtree.html) implemented as a R package^90^.

## Supporting information

Full Supplement

## Data availability

Data from this study can be found at: https://github.com/DevlinaC/FixJ_KdpE PDB structures used in Figure 1: 5XSO, [http://doi.org/10.2210/pdb5SXO/pdb], chain A and 4KFC, [http://doi.org/10.2210/pdb4KFC/pdb], chain A. All Source Data are included with the Manuscript.

## Code availability

Code used to generate the results reported in this manuscript is available at: https://github.com/DevlinaC/FixJ_KdpE

## Acknowledgements

We thank Carolyn Ott for helpful discussions and Loren Looger for critically reading this manuscript. This work utilized the NIH HPS Biowulf cluster (http://hpc.nih.gov). It was supported in part by funding from the Intramural Research Program of the National Library of Medicine, National Institutes of Health (LM202011, L.L.P.), the National Institute of General Medical Sciences, National Institutes of Health (GM118589 to L.S-K) and the W. M. Keck Foundation (L.S-K).

## Author contributions

Conceptualization: LLP, LSK
Methodology: LLP, DC, LSK, SS
Software: DC, LLP, SS
Investigation: LLP, DC, LSK, SS
Data Curation: SS, DC, LLP
Visualization: LLP, DC, SS
Writing – original draft: LLP, DC, SS
Writing – review & editing: LLP, LSK, DC, SS
Supervision: LLP, LSK
Project administration: LLP
Funding acquisition: LLP, LSK

## Competing interests

The authors declare no competing interests.

## Notes

https://github.com/DevlinaC/FixJ_KdpE

### Competing Interest Statement

The authors have declared no competing interest.

## References

1 Anfinsen, C. B. Principles that govern the folding of protein chains. Science 181, 223–230, doi:10.1126/science.181.4096.223 (1973).

2 Meinhardt, S., Manley, M. W., Jr., Parente, D. J. & Swint-Kruse, L. Rheostats and toggle switches for modulating protein function. PLoS One 8, e83502, doi:10.1371/journal.pone.0083502 (2013).

3 Markin, C. J. et al. Revealing enzyme functional architecture via high-throughput microfluidic enzyme kinetics. Science 373, doi:10.1126/science.abf8761 (2021).

4 Cole-Strauss, A. et al. Correction of the mutation responsible for sickle cell anemia by an RNA-DNA oligonucleotide. Science 273, 1386–1389, doi:10.1126/science.273.5280.1386 (1996).

5 Morral, N. et al. The origin of the major cystic fibrosis mutation (delta F508) in European populations. Nat Genet 7, 169–175, doi:10.1038/ng0694-169 (1994).

6 Muller, P. A. & Vousden, K. H. p53 mutations in cancer. Nat Cell Biol 15, 2–8, doi:10.1038/ncb2641 (2013).

7 Bai, Y. & Englander, S. W. Future directions in folding: the multi-state nature of protein structure. Proteins 24, 145–151, doi:10.1002/(SICI)1097-0134(199602)24:2<145::AID-PROT1>3.0.CO;2-I (1996).

8 Jackson, S. E. & Fersht, A. R. Folding of chymotrypsin inhibitor 2. 1. Evidence for a two-state transition. Biochemistry 30, 10428–10435, doi:10.1021/bi00107a010 (1991).

9 Orengo, C. A., Pearl, F. M. & Thornton, J. M. The CATH domain structure database. Methods Biochem Anal 44, 249–271, doi:10.1002/0471721204.ch13 (2003).

10 Andreeva, A. et al. Data growth and its impact on the SCOP database: new developments. Nucleic Acids Res 36, D419–425, doi:10.1093/nar/gkm993 (2008).

11 Greene, L. H. et al. The CATH domain structure database: new protocols and classification levels give a more comprehensive resource for exploring evolution. Nucleic Acids Res 35, D291–297, doi:10.1093/nar/gkl959 (2007).

12 Baek, M. et al. Accurate prediction of protein structures and interactions using a three-track neural network. Science 373, 871–876, doi:10.1126/science.abj8754 (2021).

13 Chowdhury, R. et al. Single-sequence protein structure prediction using a language model and deep learning. Nat Biotechnol, doi:10.1038/s41587-022-01432-w (2022).

14 Jumper, J. et al. Highly accurate protein structure prediction with AlphaFold. Nature 596, 583–589, doi:10.1038/s41586-021-03819-2 (2021).

15 Dishman, A. F. & Volkman, B. F. Unfolding the Mysteries of Protein Metamorphosis. ACS Chem Biol 13, 1438–1446, doi:10.1021/acschembio.8b00276 (2018).

16 Porter, L. L. & Looger, L. L. Extant fold-switching proteins are widespread. Proc Natl Acad Sci U S A 115, 5968–5973, doi:10.1073/pnas.1800168115 (2018).

17 Lei, X. et al. The Cancer Mutation D83V Induces an alpha-Helix to beta-Strand Conformation Switch in MEF2B. J Mol Biol 430, 1157–1172, doi:10.1016/j.jmb.2018.02.012 (2018).

18 Chang, Y. G. et al. Circadian rhythms. A protein fold switch joins the circadian oscillator to clock output in cyanobacteria. Science 349, 324–328, doi:10.1126/science.1260031 (2015).

19 Alexander, P. A., He, Y., Chen, Y., Orban, J. & Bryan, P. N. A minimal sequence code for switching protein structure and function. Proc Natl Acad Sci U S A 106, 21149–21154, doi:10.1073/pnas.0906408106 (2009).

20 He, Y., Chen, Y., Alexander, P. A., Bryan, P. N. & Orban, J. Mutational tipping points for switching protein folds and functions. Structure 20, 283–291, doi:10.1016/j.str.2011.11.018 (2012).

21 Alvarez-Carreno, C., Penev, P. I., Petrov, A. S. & Williams, L. D. Fold Evolution before LUCA: Common Ancestry of SH3 Domains and OB Domains. Mol Biol Evol 38, 5134–5143, doi:10.1093/molbev/msab240 (2021).

22 Farias-Rico, J. A., Schmidt, S. & Hocker, B. Evolutionary relationship of two ancient protein superfolds. Nat Chem Biol 10, 710–715, doi:10.1038/nchembio.1579 (2014).

23 Kumirov, V. K. et al. Multistep mutational transformation of a protein fold through structural intermediates. Protein Sci 27, 1767–1779, doi:10.1002/pro.3488 (2018).

24 Newlove, T., Konieczka, J. H. & Cordes, M. H. Secondary structure switching in Cro protein evolution. Structure 12, 569–581, doi:10.1016/j.str.2004.02.024 (2004).

25 Roessler, C. G. et al. Transitive homology-guided structural studies lead to discovery of Cro proteins with 40% sequence identity but different folds. Proc Natl Acad Sci U S A 105, 2343–2348, doi:10.1073/pnas.0711589105 (2008).

26 O’Leary, N. A. et al. Reference sequence (RefSeq) database at NCBI: current status, taxonomic expansion, and functional annotation. Nucleic Acids Res 44, D733–745, doi:10.1093/nar/gkv1189 (2016).

27 Berman, H. M. et al. The Protein Data Bank. Acta Crystallogr D Biol Crystallogr 58, 899–907, doi:10.1107/s0907444902003451 (2002).

28 Burley, S. K. et al. Protein Data Bank (PDB): The Single Global Macromolecular Structure Archive. Methods Mol Biol 1607, 627–641, doi:10.1007/978-1-4939-7000-1_26 (2017).

29 Potter, S. C. et al. HMMER web server: 2018 update. Nucleic Acids Res 46, W200–W204, doi:10.1093/nar/gky448 (2018).

30 Remmert, M., Biegert, A., Hauser, A. & Soding, J. HHblits: lightning-fast iterative protein sequence searching by HMM-HMM alignment. Nat Methods 9, 173–175, doi:10.1038/nmeth.1818 (2011).

31 Koretke, K. K., Lupas, A. N., Warren, P. V., Rosenberg, M. & Brown, J. R. Evolution of two-component signal transduction. Mol Biol Evol 17, 1956–1970, doi:10.1093/oxfordjournals.molbev.a026297 (2000).

32 Galperin, M. Y. Diversity of structure and function of response regulator output domains. Curr Opin Microbiol 13, 150–159, doi:10.1016/j.mib.2010.01.005 (2010).

33 Stock, A. M., Mottonen, J. M., Stock, J. B. & Schutt, C. E. Three-dimensional structure of CheY, the response regulator of bacterial chemotaxis. Nature 337, 745–749, doi:10.1038/337745a0 (1989).

34 Leonard, P. G., Golemi-Kotra, D. & Stock, A. M. Phosphorylation-dependent conformational changes and domain rearrangements in Staphylococcus aureus VraR activation. Proc Natl Acad Sci U S A 110, 8525–8530, doi:10.1073/pnas.1302819110 (2013).

35 Wright, G. S. A. et al. Architecture of the complete oxygen-sensing FixL-FixJ two-component signal transduction system. Sci Signal 11, doi:10.1126/scisignal.aaq0825 (2018).

36 Gao, R., Mack, T. R. & Stock, A. M. Bacterial response regulators: versatile regulatory strategies from common domains. Trends Biochem Sci 32, 225–234, doi:10.1016/j.tibs.2007.03.002 (2007).

37 Galperin, M. Y. Structural classification of bacterial response regulators: diversity of output domains and domain combinations. J Bacteriol 188, 4169–4182, doi:10.1128/JB.01887-05 (2006).

38 Aravind, L., Anantharaman, V., Balaji, S., Babu, M. M. & Iyer, L. M. The many faces of the helix-turn-helix domain: transcription regulation and beyond. FEMS Microbiol Rev 29, 231–262, doi:10.1016/j.femsre.2004.12.008 (2005).

39 Altschul, S. F. et al. Gapped BLAST and PSI-BLAST: a new generation of protein database search programs. Nucleic Acids Res 25, 3389–3402, doi:10.1093/nar/25.17.3389 (1997).

40 Kim, A. K., Looger, L. L. & Porter, L. L. A high-throughput predictive method for sequence-similar fold switchers. Biopolymers, e23416, doi:10.1002/bip.23416 (2021).

41 Gonzalez, M. W. & Pearson, W. R. Homologous over-extension: a challenge for iterative similarity searches. Nucleic Acids Res 38, 2177–2189, doi:10.1093/nar/gkp1219 (2010).

42 Belogurov, G. A. et al. Structural basis for converting a general transcription factor into an operon-specific virulence regulator. Mol Cell 26, 117–129, doi:10.1016/j.molcel.2007.02.021 (2007).

43 Eddy, S. R. A new generation of homology search tools based on probabilistic inference. Genome Inform 23, 205–211 (2009).

44 Sievers, F. et al. Fast, scalable generation of high-quality protein multiple sequence alignments using Clustal Omega. Mol Syst Biol 7, 539, doi:10.1038/msb.2011.75 (2011).

45 Edgar, R. C. MUSCLE: multiple sequence alignment with high accuracy and high throughput. Nucleic Acids Res 32, 1792–1797, doi:10.1093/nar/gkh340 (2004).

46 Shimodaira, H. An approximately unbiased test of phylogenetic tree selection. Syst Biol 51, 492–508, doi:10.1080/10635150290069913 (2002).

47 Chakravarty, D. & Porter, L. L. AlphaFold2 fails to predict protein fold switching. Protein Sci 31, e4353, doi:10.1002/pro.4353 (2022).

48 Porter, L. L. et al. Many dissimilar NusG protein domains switch between alpha-helix and betasheet folds. Nat Commun 13, 3802, doi:10.1038/s41467-022-31532-9 (2022).

49 Rost, B. Twilight zone of protein sequence alignments. Protein Eng 12, 85–94, doi:10.1093/protein/12.2.85 (1999).

50 Bateman, A. et al. The Pfam protein families database. Nucleic Acids Res 32, D138–141, doi:10.1093/nar/gkh121 (2004).

51 Liberles, D. A. et al. The interface of protein structure, protein biophysics, and molecular evolution. Protein Sci 21, 769–785, doi:10.1002/pro.2071 (2012).

52 Yadid, I., Kirshenbaum, N., Sharon, M., Dym, O. & Tawfik, D. S. Metamorphic proteins mediate evolutionary transitions of structure. Proc Natl Acad Sci U S A 107, 7287–7292, doi:10.1073/pnas.0912616107 (2010).

53 Alva, V., Soding, J. & Lupas, A. N. A vocabulary of ancient peptides at the origin of folded proteins. Elife 4, e09410, doi:10.7554/eLife.09410 (2015).

54 Kolodny, R., Nepomnyachiy, S., Tawfik, D. S. & Ben-Tal, N. Bridging Themes: Short Protein Segments Found in Different Architectures. Mol Biol Evol 38, 2191–2208, doi:10.1093/molbev/msab017 (2021).

55 Nepomnyachiy, S., Ben-Tal, N. & Kolodny, R. Complex evolutionary footprints revealed in an analysis of reused protein segments of diverse lengths. Proc Natl Acad Sci U S A 114, 11703–11708, doi:10.1073/pnas.1707642114 (2017).

56 Qiu, K., Ben-Tal, N. & Kolodny, R. Similar protein segments shared between domains of different evolutionary lineages. Protein Sci 31, e4407, doi:10.1002/pro.4407 (2022).

57 Alexander, P. A., He, Y., Chen, Y., Orban, J. & Bryan, P. N. The design and characterization of two proteins with 88% sequence identity but different structure and function. Proc Natl Acad Sci U S A 104, 11963–11968, doi:10.1073/pnas.0700922104 (2007).

58 Dishman, A. F. et al. Evolution of fold switching in a metamorphic protein. Science 371, 86–90, doi:10.1126/science.abd8700 (2021).

59 Grishin, N. V. Fold change in evolution of protein structures. J Struct Biol 134, 167–185, doi:10.1006/jsbi.2001.4335 (2001).

60 Ovchinnikov, S. et al. Protein structure determination using metagenome sequence data. Science 355, 294–298, doi:10.1126/science.aah4043 (2017).

61 Kabsch, W. & Sander, C. Dictionary of protein secondary structure: pattern recognition of hydrogen-bonded and geometrical features. Biopolymers 22, 2577–2637, doi:10.1002/bip.360221211 (1983).

62 Cheng, H. et al. ECOD: an evolutionary classification of protein domains. PLoS Comput Biol 10, e1003926, doi:10.1371/journal.pcbi.1003926 (2014).

63 Wang, Y., Wu, H. & Cai, Y. A benchmark study of sequence alignment methods for protein clustering. BMC bioinformatics 19, 529, doi:10.1186/s12859-018-2524-4 (2018).

64 Cock, P. J. et al. Biopython: freely available Python tools for computational molecular biology and bioinformatics. Bioinformatics 25, 1422–1423, doi:10.1093/bioinformatics/btp163 (2009).

65 Pei, J. & Grishin, N. V. PROMALS: towards accurate multiple sequence alignments of distantly related proteins. Bioinformatics 23, 802–808, doi:10.1093/bioinformatics/btm017 (2007).

66 Ashkenazy, H. et al. ConSurf 2016: an improved methodology to estimate and visualize evolutionary conservation in macromolecules. Nucleic Acids Res 44, W344–350, doi:10.1093/nar/gkw408 (2016).

67 Parente, D. J., Ray, J. C. J. & Swint-Kruse, L. Amino acid positions subject to multiple coevolutionary constraints can be robustly identified by their eigenvector network centrality scores. Proteins 83, 2293–2306, doi:10.1002/prot.24948 (2015).

68 Bolten, E., Schliep, A., Schneckener, S., Schomburg, D. & Schrader, R. Clustering protein sequences--structure prediction by transitive homology. Bioinformatics 17, 935–941, doi:10.1093/bioinformatics/17.10.935 (2001).

69 Gerstein, M. Measurement of the effectiveness of transitive sequence comparison, through a third ‘intermediate’ sequence. Bioinformatics 14, 707–714, doi:10.1093/bioinformatics/14.8.707 (1998).

70 Fu, L., Niu, B., Zhu, Z., Wu, S. & Li, W. CD-HIT: accelerated for clustering the next-generation sequencing data. Bioinformatics 28, 3150–3152, doi:10.1093/bioinformatics/bts565 (2012).

71 Mayrose, I., Graur, D., Ben-Tal, N. & Pupko, T. Comparison of site-specific rate-inference methods for protein sequences: empirical Bayesian methods are superior. Mol Biol Evol 21, 1781–1791, doi:10.1093/molbev/msh194 (2004).

72 Price, M. N., Dehal, P. S. & Arkin, A. P. FastTree: computing large minimum evolution trees with profiles instead of a distance matrix. Mol Biol Evol 26, 1641–1650, doi:10.1093/molbev/msp077 (2009).

73 Price, M. N., Dehal, P. S. & Arkin, A. P. FastTree 2--approximately maximum-likelihood trees for large alignments. PLoS One 5, e9490, doi:10.1371/journal.pone.0009490 (2010).

74 Jones, D. T., Taylor, W. R. & Thornton, J. M. The rapid generation of mutation data matrices from protein sequences. Comput Appl Biosci 8, 275–282, doi:10.1093/bioinformatics/8.3.275 (1992).

75 Stamatakis, A. in Proceedings 20th IEEE international parallel & distributed processing symposium. 8 pp. (IEEE).

76 Hoang, D. T., Chernomor, O., von Haeseler, A., Minh, B. Q. & Vinh, L. S. UFBoot2: Improving the Ultrafast Bootstrap Approximation. Mol Biol Evol 35, 518–522, doi:10.1093/molbev/msx281 (2018).

77 Nguyen, L. T., Schmidt, H. A., von Haeseler, A. & Minh, B. Q. IQ-TREE: a fast and effective stochastic algorithm for estimating maximum-likelihood phylogenies. Mol Biol Evol 32, 268–274, doi:10.1093/molbev/msu300 (2015).

78 Kalyaanamoorthy, S., Minh, B. Q., Wong, T. K. F., von Haeseler, A. & Jermiin, L. S. ModelFinder: fast model selection for accurate phylogenetic estimates. Nat Methods 14, 587–589, doi:10.1038/nmeth.4285 (2017).

79 Naser-Khdour, S., Quang Minh, B. & Lanfear, R. Assessing Confidence in Root Placement on Phylogenies: An Empirical Study Using Nonreversible Models for Mammals. Syst Biol 71, 959–972, doi:10.1093/sysbio/syab067 (2022).

80 Kishino, H., Miyata, T. & Hasegawa, M. Maximum likelihood inference of protein phylogeny and the origin of chloroplasts. Journal of Molecular Evolution 31, 151–160 (1990).

81 Kishino, H. & Hasegawa, M. Evaluation of the maximum likelihood estimate of the evolutionary tree topologies from DNA sequence data, and the branching order in hominoidea. J Mol Evol 29, 170–179, doi:10.1007/BF02100115 (1989).

82 Shimodaira, H. & Hasegawa, M. Multiple comparisons of log-likelihoods with applications to phylogenetic inference. Molecular biology and evolution 16, 1114 (1999).

83 Strimmer, K. & Rambaut, A. Inferring confidence sets of possibly misspecified gene trees. Proc Biol Sci 269, 137–142, doi:10.1098/rspb.2001.1862 (2002).

84 Yang, Z., Kumar, S. & Nei, M. A new method of inference of ancestral nucleotide and amino acid sequences. Genetics 141, 1641–1650, doi:10.1093/genetics/141.4.1641 (1995).

85 Varadi, M. et al. AlphaFold Protein Structure Database: massively expanding the structural coverage of protein-sequence space with high-accuracy models. Nucleic Acids Research 50, D439–D444, doi:10.1093/nar/gkab1061 (2021).

86 Swint-Kruse, L. & Brown, C. S. Resmap: automated representation of macromolecular interfaces as two-dimensional networks. Bioinformatics 21, 3327–3328, doi:10.1093/bioinformatics/bti511 (2005).

87 The PyMOL Molecular Graphics System, Version 2.0 Schrödinger, LLC.

88 Hunter, J. D. Matplotlib: A 2D graphics environment. Comput Sci Eng 9, 90–95 (2007).

89 Waskom, M. L. seaborn: statistical data visualization. Journal of Open Source Software 6, doi:10.21105/joss.03021 (2021).

90 Yu, G., Smith, D. K., Zhu, H., Guan, Y. & Lam, T. T. Y. ggtree: an R package for visualization and annotation of phylogenetic trees with their covariates and other associated data. Methods in Ecology and Evolution 8, 28–36 (2017).

