## Supplementary material for "Identification of a covert evolutionary pathway between two protein folds": Full Supplement

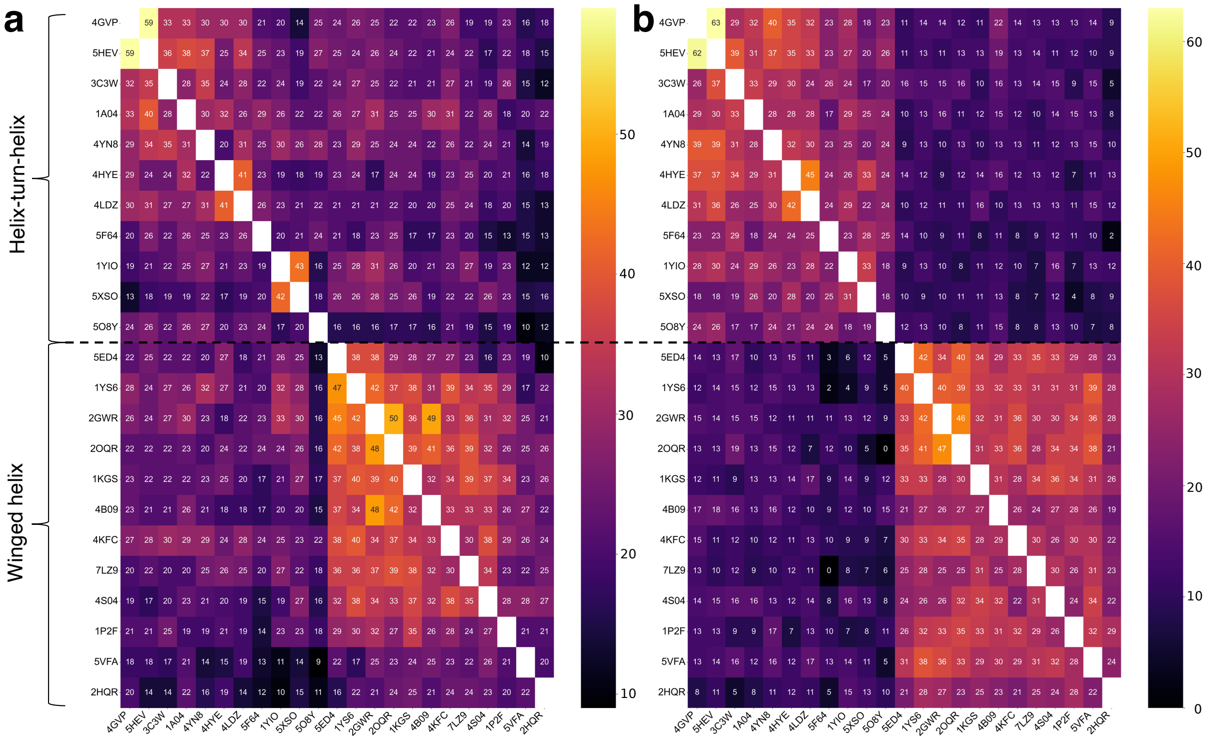


*Figure S1. Response regulator subfamilies cluster by CTD architecture (left labels). Sequence comparisons between response regulators with experimentally determined structures were performed with jackhmmer[1]. PDB codes are indicated on the left and bottom axes of each panel. In these sequence identity matrices, structures with $HTH_4$ CTDS cluster in the upper left, whereas those with wH CTDs cluster in the lower right. The color bar indicates high (yellow) to low (black) sequence identity. (a) Sequence identity matrix of the NTDs lacks a strong distinction between response regulators with CTDs from the two fold families. (b). By contrast, the sequence identity matrix of the CTDs shows two distinct clusters that correspond to the different CTD architectures. Black dotted line separates $HTH_4$ structures (above) and wH structures (below).*

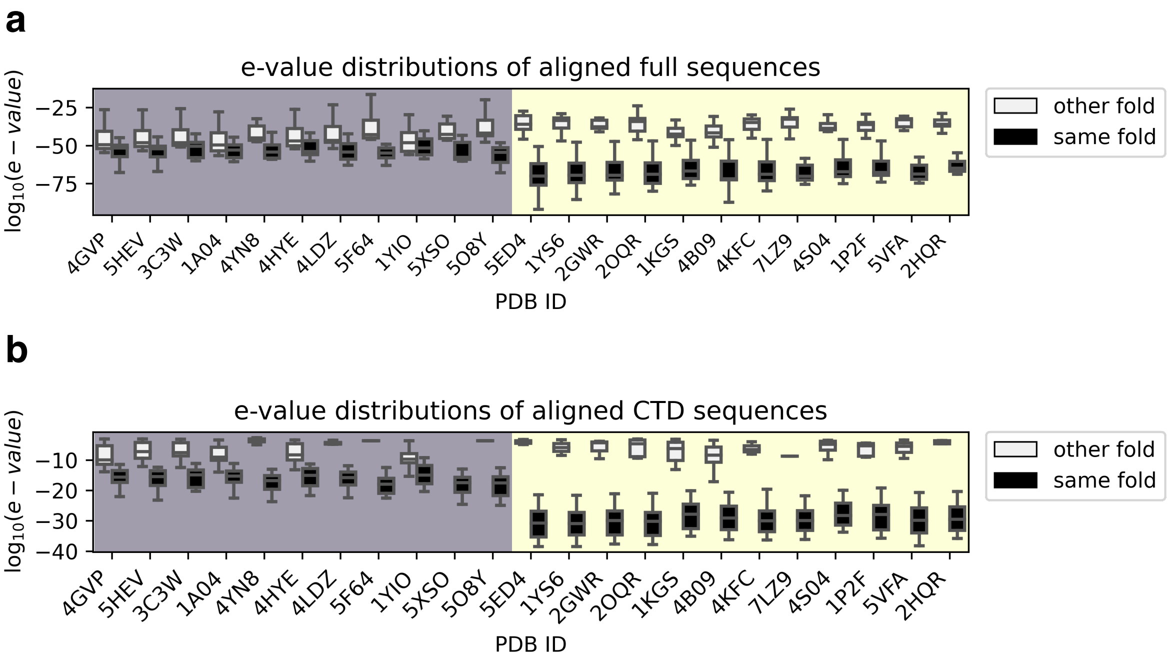


*Figure S2. Box-and-whisker plots of $log_{10}$(e-values) from jackhmmer searches of the 23 response regulators in the PDB. Sequence comparisons were made between (a) full-length sequences and (b) CTDs with (gray boxes) structures with the same fold and (white boxes) those with the other fold. The distributions of each HTH/wH box were derived from n=1-26 jackhmmer e-values (Source Data); each box bounds the interquartile range (IQR) of the data (first quartile, Q1 through third quartile, Q3); medians of each distribution are gray lines within each black box; lower whisker is the lowest datum above Q1-1.5*IQR; upper whisker is the highest datum below Q3 + 1.5*IQR. E-values < 5e-02 are considered significant. Note that the e-values of (b) are expected to be lower than those of (a) because the sequences are shorter.*

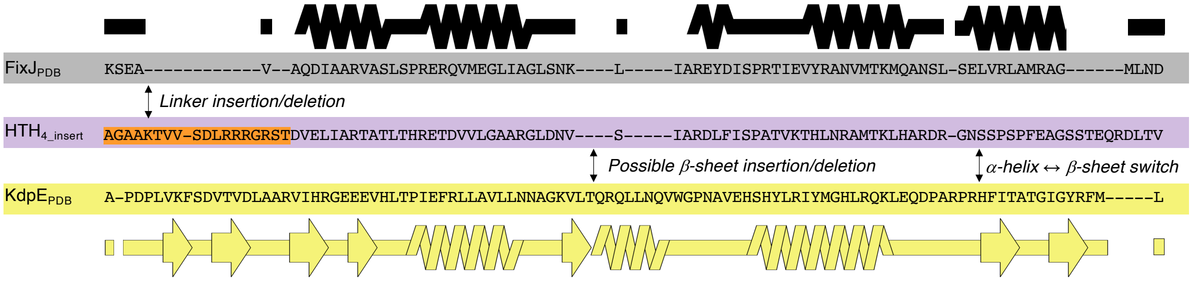


*Figure S3. MUSCLE[2] alignment of 3,205 sequences also suggests a mutationally-driven secondary structure conversion. Secondary structure diagrams were generated from the structures of FixJ$_{PDB}$ (black) and KdpE$_{PDB}$ (yellow). The features of this diagram are analogous to those described in Fig 3a. Spaces between sequences show important changes: (1) orange linker insertion/deletion (2) fold conversion and possible β-sheet insertion/deletion.*

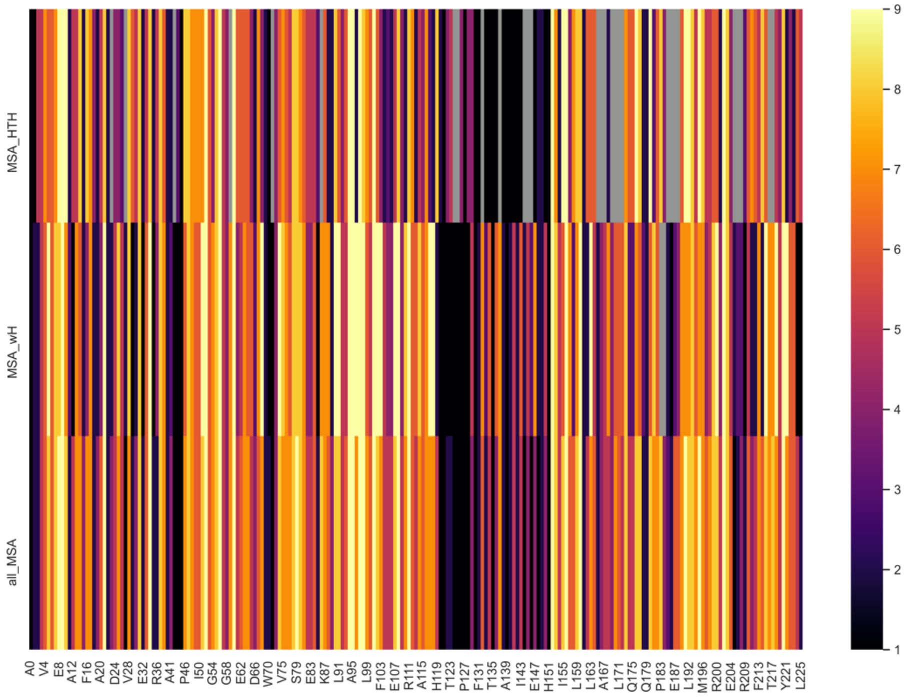


*Figure S4. Evolutionary rates of only $HTH_4$ (664, top), only wH (2541, middle), and all 3205 sequences of the cross-family MSA (bottom) of Figure 3b. These rates were calculated using ConSurf[3] to analyze the Clustal Omega[4] alignment and the consensus tree. Position numbers at the bottom of the plot correspond to those of $KdpE_{PDB}$ (PBD ID: 4KFC, chain A). The conservation scale (color bar) ranges from 1 (not conserved, black) to 9 (highly conserved, yellow); gray represents alignment gaps. The greater number of wH sequences (Figure 3) probably biases the results of the "all_MSA" calculation. Conservation patterns in the N-terminal domains (positions 0-119) of all 3 sets of sequences show similar rates in similar positions. In both subfamilies, the inter-domain linker shows more rapid evolution (darker colors) than either the NTD or CTD. By contrast, conservation patterns in the C-terminal domains differ. Most notably, several positions in the wing of the wH sequences (positions 213-225) are highly conserved (score of 9), whereas corresponding positions in the $HTH_4$ evolve more rapidly (scores of 1-6).*

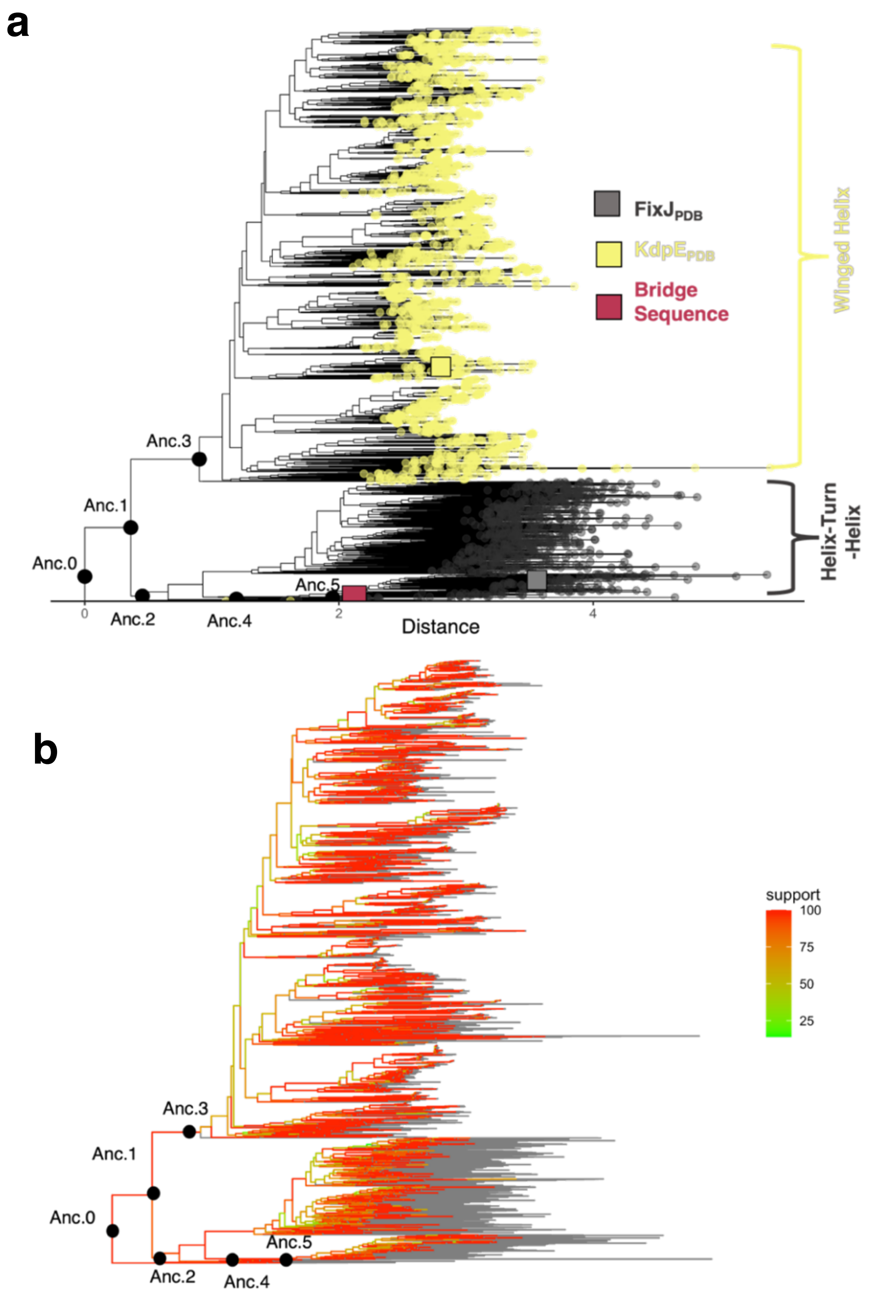


*Figures S5. (a) Unrooted consensus tree built after ultrafast bootstrapping using the maximum likelihood (ML) tree (Figure 3) as input. The Robinson-Foulds distance (which denotes the number of splits needed to convert one tree to another) between this tree and the ML tree in Figure 3 is 127, indicating 98% similarity between the two trees, and confirming the accuracy of the tree shown in Figure 3b. The bridge sequences in this unrooted consensus tree can also lie between the fold families if all branches connected at Anc. 2 are flipped by 180°. Thus, we show the ML tree in Figure 3b to highlight that the branch containing the bridge sequences can adjoin both fold families. Nodes annotated as helix-turn-helix (HTH) and winged-helix (wH) are colored yellow and gray, respectively. This consensus tree was used for ancestral reconstruction (Figure 4); ancestral nodes are marked as black circles. (b) Bootstrapping support shown on the consensus tree after performing ultrafast bootstrapping for 1000 replicates at 500 iterations. Nodes are color-coded by the level of support shown in the color bar. Most branches are well supported with values > 80%. Importantly, all ancestral nodes (black circles) are supported with values > 95%. Locations of the unsupported gray branches are ambiguous.*

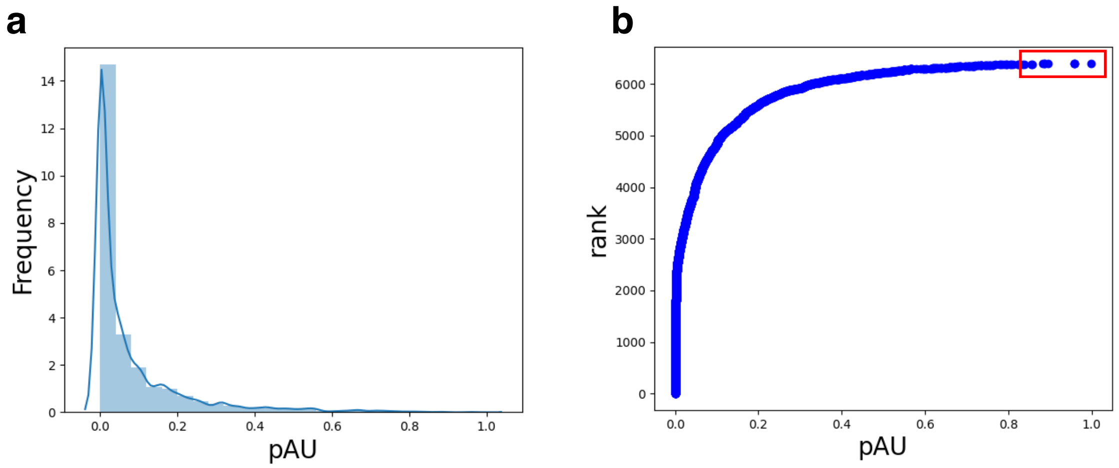


*Figures S6. Frequencies (a) and ranks (b) of probable tree rootings from the approximately unbiased test (pAU) of the tree shown in Figure 3b and Figure S5. Although most rootings were highly improbable, 18of 6393 had a pAU ≥0.8 (b, red box), suggesting likely rootings.*

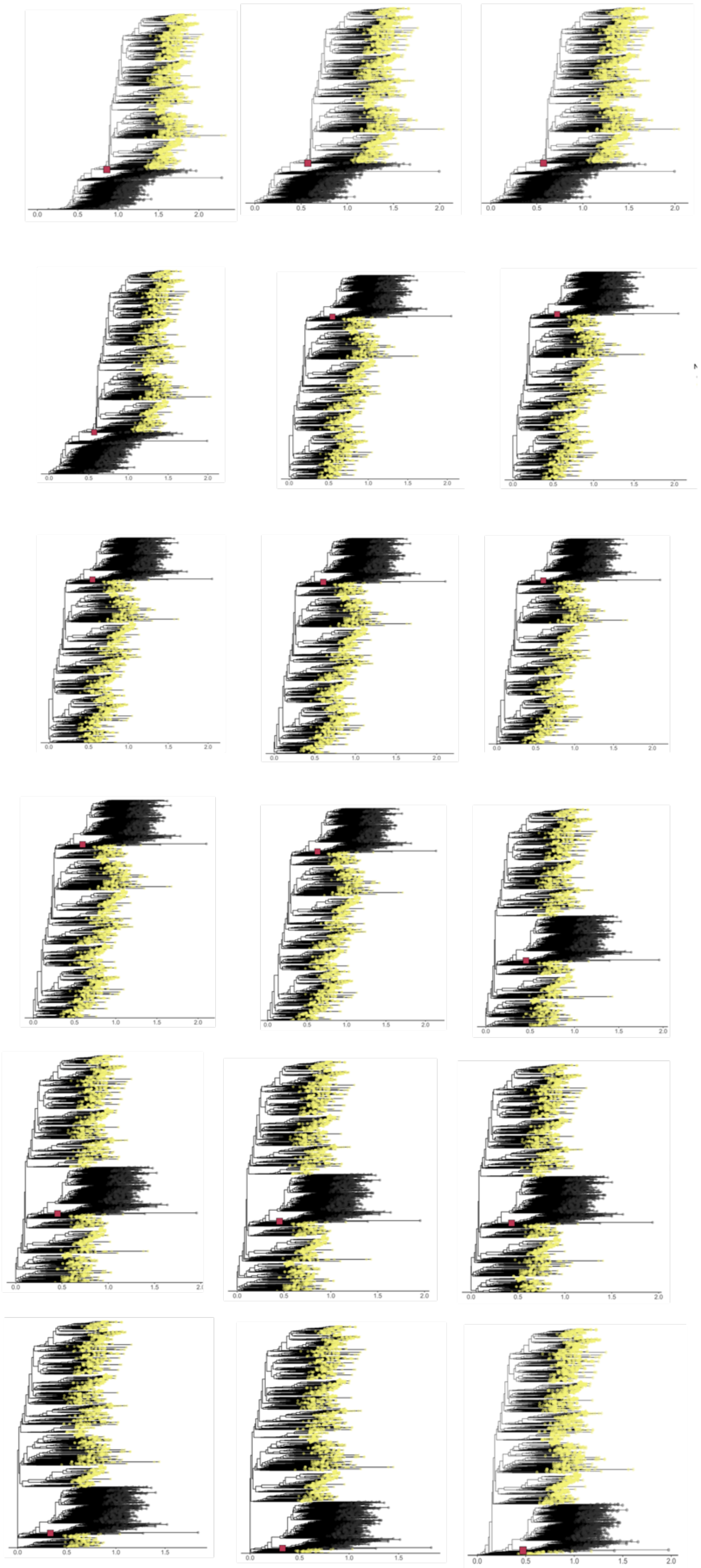

*Figure S7. Ancestors to the branch containing the 12 bridge sequences may have served as the evolutionary intermediates between helix-turn-helix and winged helix folds. The branches with bridge sequences (pink square) adjoin branches with helix-turn-helix (gray) and winged helix (yellow) sequences in all 18 phylogenetic trees with most probable rootings.*

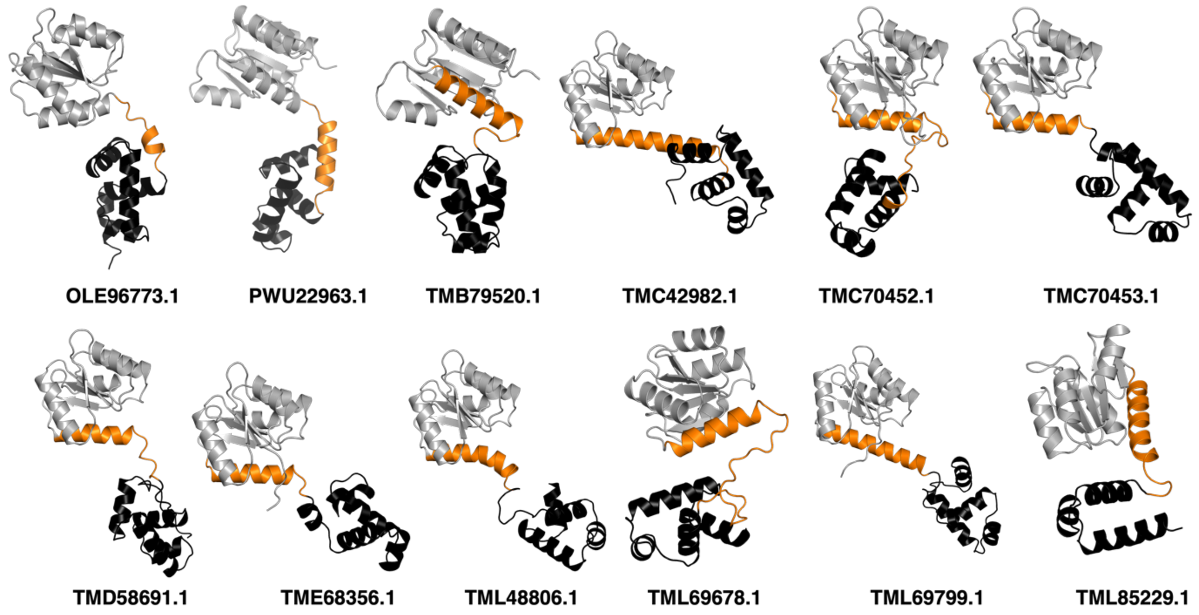


*Figure S8. AlphaFold2 structure predictions for the 12 bridge sequences. In all 12 cases, the C-terminal DNA-binding domains (black) are predicted to assume helix-turn-helix folds, suggesting that evolved fold switching may have occurred in ancestral sequences. The N-terminal receiver domains are gray, and the linkers are orange. NCBI Protein database accession codes are shown below each model; their sequences are reported in* ***Table S4****.*

*Table S1. PSI-BLAST[5] searches of $\text{FixJ}_{PDB}$ sequence (query, PDB ID: 5XSO, chain A) against all the sequences found in the PDB identified dozens of evolutionarily-related sequences, the top 20 of which are shown below. The three columns show the PDB ID, e-value, and alignment, respectively. The CTD of $\text{FixJ}_{PDB}$ assumes a helix-turn-helix fold. White rows indicate alignments with other response regulators with CTD helix-turn-helix folds. Gray rows indicate alignments with response regulators with CTD winged helix folds.*

| PDB ID | e-value | Alignment |
|---|---|---|
| 5XSO | 7e-55 | Query 1 MTTKGHIYVIDDDAAMRDSLNFLLDSAGFGVTLFDDAQAFLDALPGLSFGCVVSDVRMPG 60<br> MTTKGHIYVIDDDAAMRDSLNFLLDSAGFGVTLFDDAQAFLDALPGLSFGCVVSDVRMPG<br>Sbjct 11 MTTKGHIYVIDDDAAMRDSLNFLLDSAGFGVTLFDDAQAFLDALPGLSFGCVVSDVRMPG 70<br><br>Query 61 LDGIELLKRMKAQQSPFPILIMTGHGDVPLAVEAMKLGAVDFLEKPFEDDRLTAMIESAI 120<br> LDGIELLKRMKAQQSPFPILIMTGHGDVPLAVEAMKLGAVDFLEKPFEDDRLTAMIESAI<br>Sbjct 71 LDGIELLKRMKAQQSPFPILIMTGHGDVPLAVEAMKLGAVDFLEKPFEDDRLTAMIESAI 130<br><br>Query 121 RQAEPAAKSEAVAQDIAARVASLSPRERQVMEGLIAGLSNKLIAREYDISPRTIEVYRAN 180<br> RQAEPAAKSEAVAQDIAARVASLSPRERQVMEGLIAGLSNKLIAREYDISPRTIEVYRAN<br>Sbjct 131 RQAEPAAKSEAVAQDIAARVASLSPRERQVMEGLIAGLSNKLIAREYDISPRTIEVYRAN 190<br><br>Query 181 VMTKMQANSLSELVRLAMRAGMLND 205<br> VMTKMQANSLSELVRLAMRAGMLND<br>Sbjct 191 VMTKMQANSLSELVRLAMRAGMLND 215 |
| 1YIO | 2e-48 | Query 1 MTTKGHIYVIDDDAAMRDSLNFLLDSAGFGVTLFDDAQAFLDALPGLSFGCVVSDVRMPG 60<br> MT K ++V+DDD ++R+ L LL SAGF V FD A FL+ GC+V D+RMPG<br>Sbjct 1 MTAKPTVFVVDDDMSVREGLRNLLRSAGFEVETFDCASTFLEHRRPEQHGCLVLDMRMPG 60<br><br>Query 61 LDGIELLKRMKAQQSPFPILIMTGHGDVPLAVEAMKLGAVDFLEKPFEDDRLTAMIESAI 120<br> + GIEL +++ A PI+ +T HGD+P+ V AMK GA++FL KPFE+ L IE +<br>Sbjct 61 MSGIELQEQLTAISDGIPIVFITAHGDIPMTVRAMKAGAIEFLPKPFEEQALLDAIEQGL 120<br><br>Query 121 RQAEPAAKSEAVAQDIAARVASLSPRERQVMEGLIAGLSNKLIAREYDISPRTIEVYRAN 180<br> + ++ + +SL+ RE+QV++ I GL NK IA E I+ T++V+R N<br>Sbjct 121 QLNAERRQARETQDQLEQLFSSLTGREQQVLQLTIRGLMNKQIAGELGIAEVTVKVHRHN 180<br><br>Query 181 VMTKMQANSLSELVRLAMR 199<br> +M K+ SL+ LV L +<br>Sbjct 181 IMQKLNVRSLANLVHLVEK 199 |

| | | |
|---|---|---|
| 4HYE | 1e-42 | Query 5 GHIYVIDDDAAMRDSLNFLL--DSAGFGVTLFDDAQAFLDALPGLSFGCVVSDVRMPGLD 62<br> + V +D + +RD++ LL V + Q + L S + DV MP<br>Sbjct 22 MKVLVAEDQSMLRDAMCQLLTLQPDVESVLQAKNGQEAIQLLEKESVDIAILDVEMPVKT 81<br><br>Query 63 GIELLKRMKAQQSPFPILIMTGHGDVPLAVEAMKLGAVDFLEKPFEDDRLTAMIESAIRQ 122<br> G+E+L+ +++++ ++++T A+K G ++ K L + + +<br>Sbjct 82 GLEVLEWIRSEKLETKVVVVTTFKRAGYFERAVKAGVDAYVLKERSIADLMQTLHTVLEG 141<br><br>Query 123 AEPAAKSEAVAQDIAARVASLSPRERQVMEGLIAGLSNKLIAREYDISPRTIEVYRANVM 182<br> + + + + + R L+ +E V++G+ GLSN+ IA + +S TI Y N++<br>Sbjct 142 RKEYSPE--LMEMVMTRPNPLTEQEIAVLKGIARGLSNQEIADQLYLSNGTIRNYVTNIL 199<br><br>Query 183 TKMQANSLSELVRLAMRAGML 203<br> +K+ A + +E +A +G L<br>Sbjct 200 SKLDAGNRTEAANIAKESGWL 220 |
| 4ZMR | 2e-42 | Query 5 GHIYVIDDDAAMRDSLNFLL--DSAGFGVTLFDDAQAFLDALPGLSFGCVVSDVRMPGLD 62<br> + V +D + +RD++ LL V + Q + L S + DV MP<br>Sbjct 9 MKVLVAEDQSMLRDAMCQLLTLQPDVESVLQAKNGQEAIQLLEKESVDIAILDVEMPVKT 68<br><br>Query 63 GIELLKRMKAQQSPFPILIMTGHGDVPLAVEAMKLGAVDFLEKPFEDDRLTAMIESAIRQ 122<br> G+E+L+ +++++ ++++T A+K G ++ K L + + +<br>Sbjct 69 GLEVLEWIRSEKLETKVVVVTTFKRAGYFERAVKAGVDAYVLKERSIADLMQTLHTVLEG 128<br><br>Query 123 AEPAAKSEAVAQDIAARVASLSPRERQVMEGLIAGLSNKLIAREYDISPRTIEVYRANVM 182<br> + + + + + R L+ +E V++G+ GLSN+ IA + +S TI Y N++<br>Sbjct 129 RKEYSPE--LMEMVMTRPNPLTEQEIAVLKGIARGLSNQEIADQLYLSNGTIRNYVTNIL 186<br><br>Query 183 TKMQANSLSELVRLAMRAGML 203<br> +K+ A + +E +A +G L<br>Sbjct 187 SKLDAGNRTEAANIAKESGWL 207 |

| | | |
|---|---|---|
| 5ED4 | 4e-41 | Query 2 TTKGHIYVIDDDAAMRDSLNFLLDSAGFGVTLFDDAQAFLDALPGLSFGCVVSDVRMPGL 61<br>T + + V+DD+A + + L+ L GF V + LD V+ DV MPG+<br>Sbjct 21 TPEARVLVVDDEANIVELLSVSLKFQGFEVYTATNGAQALDRARETRPDAVILDVMMPGM 80<br><br>Query 62 DGIELLKRMKAQQSPFPILIMTGHGDVPLAVEAMKLGAVDFLEKPFEDDRLTAMIESAIR 121<br>DG +L+R++A P L +T + + + LG D++ KPF + + A + +R<br>Sbjct 81 DGFGVLRRLRADGIDAPALFLTARDSLQDKIAGLTLGGDDYVTKPFSLEEVVARLRVILR 140<br><br>Query 122 QA----EPAAKSEAVAQDIAARVA-----------SLSPRERQVMEGLIAGLSNK----- 161<br>+A + DI SLSP E ++ +<br>Sbjct 141 RAGKGNKEPRNVRLTFADIELDEETHEVWKAGQPVSLSPTEFTLLRYFVINAGTVLSKPK 200<br><br>Query 162 ----LIAREYDISPRTIEVYRANVMTKMQANSLSELVRLAMRAGML 203<br>+ ++ +E Y + + K+ L+ G +<br>Sbjct 201 ILDHVWRYDFGGDVNVVESYVSYLRRKIDTGEK-RLLHTLRGVGYV 245 |
| 5HEV | 9e-41 | Query 4 KGHIYVIDDDAAMRDSLNFLLD-SAGFGVT-LFDDAQAFLDALPGLSFGCVVSDVRMPGL 61<br>+ ++DD +R ++ L V ++ + + L ++ D+ M +<br>Sbjct 1 MIKVLLVDDHEMVRLGVSSYLSIQEDIEVIGEAENGRQGYEKAMALRPDVILMDLVMEEM 60<br><br>Query 62 DGIELLKRMKAQQSPFPILIMTGHGDVPLAVEAMKLGAVDFLEKPFEDDRLTAMIESAIR 121<br>DGIE K + I+I+T D A++ GA +L K + I + R<br>Sbjct 61 DGIESTKAILKDWPKAKIIIVTSFIDDEKVYPAIEAGAAGYLLKTSTAHEIADAIRATQR 120<br><br>Query 122 --QAEPAAKSEAVAQDIAARVAS-----LSPRERQVMEGLIAGLSNKLIAREYDISPRTI 174<br>+ + + + ++ R L+ RE +++ + G SN+ IA E I+ +T+<br>Sbjct 121 GERVLEPEVTTKMMEKMSRRNDPVLHEELTNRENEILMLISEGKSNQEIADELFITLKTV 180<br><br>Query 175 EVYRANVMTKMQANSLSELVRLAMRAGML 203<br>+ + +N++ K++ ++ A + G++<br>Sbjct 181 KTHVSNILAKLEVEDRTQAAIYAFKHGLV 209 |

| | | |
|---|---|---|
| 1KGS | 3e-40 | Query 4 KGHIYVIDDDAAMRDSLNFLLDSAGFGVTLFDDAQAFLDALPGLSFGCVVSDVRMPGLDG 63<br> + V++D+ + D + L F V + D + F V+ D+ +P DG<br>Sbjct 2 NVRVLVVEDERDLADLITEALKKEXFTVDVCYDGEEGXYXALNEPFDVVILDIXLPVHDG 61<br><br>Query 64 IELLKRMKAQQSPFPILIMTGHGDVPLAVEAMKLGAVDFLEKPFEDDRLTAMIESAIRQA 123<br> E+LK + P+L +T DV V+ + GA D+L KPF+ L A + + IR+<br>Sbjct 62 WEILKSXRESGVNTPVLXLTALSDVEYRVKGLNXGADDYLPKPFDLRELIARVRALIRRK 121<br><br>Query 124 EPAAKSEAVAQDI-----------AARVASLSPRERQVMEGLIAGLSNKLIAREY----- 167<br> + ++ V D+ ++ L+ +E Q++E L+ + + E<br>Sbjct 122 SESKSTKLVCGDLILDTATKKAYRGSKEIDLTKKEYQILEYLVXNKNRVVTKEELQEHLW 181<br><br>Query 168 ----DISPRTIEVYRANVMTKMQANSLSELVRLAMRAGML 203<br> ++ + + N+ K+ +++ G +<br>Sbjct 182 SFDDEVFSDVLRSHIKNLRKKVDKGFKKKIIHTVRGIGYV 221 |
| 4GVP | 4e-40 | Query 4 KGHIYVIDDDAAMRDSLNFLLD-SAGFGVT-LFDDAQAFLDALPGLSFGCVVSDVRMPGL 61<br> + +DD +R ++ L + V + + L ++ D+ M +<br>Sbjct 1 TIKVLFVDDHEMVRIGISSYLSTQSDIEVVGEGASGKEAIAKAHELKPDLILMDLLMEDM 60<br><br>Query 62 DGIELLKRMKAQQSPFPILIMTGHGDVPLAVEAMKLGAVDFLEKPFEDDRLTAMIESAIR 121<br> DG+E ++K +L++T + A+ G ++ K + + R<br>Sbjct 61 DGVEATTQIKKDLPQIKVLMLTSFIEDKEVYRALDAGVDSYILKTTSAKDIADAVRKTSR 120<br><br>Query 122 QAEPAAKSEAV-----AQDIAARVASLSPRERQVMEGLIAGLSNKLIAREYDISPRTIEV 176<br> V + A L+ RE +++ + G SN+ IA I+ +T++<br>Sbjct 121 GESVFEPEVLVKMRNRMKKRAELYEMLTEREMEILLLIAKGYSNQEIASASHITIKTVKT 180<br><br>Query 177 YRANVMTKMQANSLSELVRLAMRAGML 203<br> + +N+++K++ ++ V A + ++<br>Sbjct 181 HVSNILSKLEVQDRTQAVIYAFQHNLI 207 |

| | | |
|---|---|---|
| 4LDZ | 1e-39 | Query 3 TKGHIYVIDDDAAMRDSLNFLLD-SAGFGVTL-FDDAQAFLDALPGLSFGCVVSDVRMPG 60<br>+ I++ +D + +L LL+ V Q +D + + D+ MPG<br>Sbjct 4 SMISIFIAEDQQMLLGALGSLLNLEDDMEVVGKGTTGQDAVDFVKKRQPDVCIMDIEMPG 63<br><br>Query 61 LDGIELLKRMKAQQSPFPILIMTGHGDVPLAVEAMKLGAVDFLEKPFEDDRLTAMIESAI 120<br>G+E + +K + I+I+T A+K G +L K +L I S +<br>Sbjct 64 KTGLEAAEELK--DTGCKIIILTTFARPGYFQRAIKAGVKGYLLKDSPSEELANAIRSVM 121<br><br>Query 121 RQAEPAAKSEAVAQDIAARVASLSPRERQVMEGLIAGLSNKLIAREYDISPRTIEVYRAN 180<br>A + +D+ + L+ RE++V+E + G + K IA+E I T+ Y +<br>Sbjct 122 NGKRIYAPE--LMEDLYSEANPLTDREKEVLELVADGKNTKEIAQELSIKSGTVRNYISM 179<br><br>Query 181 VMTKMQANSLSELVRLAMRAGM 202<br>++ K++ + E + + G<br>Sbjct 180 ILEKLEVKNRIEAITRSKEKGW 201 |
| 4KFC | 6e-39 | Query 3 TKGHIYVIDDDAAMRDSLNFLLDSAGFGVTLFDDAQAFLDALPGLSFGCVVSDVRMPGLD 62<br>++ +++D+ A+R L L+ G V + Q L ++ D+ +P D<br>Sbjct 2 AMANVLIVEDEQAIRRFLRTALEGDGMRVFEAETLQRGLLEAATRKPDLIILDLGLPDGD 61<br><br>Query 63 GIELLKRMKAQQSPFPILIMTGHGDVPLAVEAMKLGAVDFLEKPFEDDRLTAMIESAIRQ 122<br>GIE ++ ++ S P+++++ + + A+ GA D+L KPF L A + A+R+<br>Sbjct 62 GIEFIRDLRQ-WSAVPVIVLSARSEESDKIAALDAGADDYLSKPFGIGELQARLRVALRR 120<br><br>Query 123 A------EPAAKSEAVAQDIAAR-------VASLSPRERQVMEGLIAGLSNKLIAREYD- 168<br>+P K V D+AAR L+P E +++ L+ L R+<br>Sbjct 121 HSATTAPDPLVKFSDVTVDLAARVIHRGEEEVHLTPIEFRLLAVLLNNAGKVLTQRQLLN 180<br><br>Query 169 --------ISPRTIEVYRANVMTKMQANS-LSELVRLAMRAGM 202<br>+ +Y ++ K++ + A G<br>Sbjct 181 QVWGPNAVEHSHYLRIYMGHLRQKLEQDPARPRHFITATGIGY 223 |

<table>
<tr>
<td>3R0J</td>
<td>1e-38</td>
<td><pre>
Query  2    TTKGHIYVIDDDAAMRDSLNFLLDSAGFGVTLFDDAQAFLDALPGLSFGCVVSDVRMPGL  61
            T +  + V+DD+A + + L+  L   GF V    +    LD         V+ DV  PG
Sbjct  21   TPEARVLVVDDEANIVELLSVSLKFQGFEVYTATNGAQALDRARETRPDAVILDVXXPGX  80

Query  62   DGIELLKRMKAQQSPFPILIMTGHGDVPLAVEAMKLGAVDFLEKPFEDDRLTAMIESAIR  121
            DG  +L+R++A     P L +T    +   +  + LG  D++ KPF  + + A +   +R
Sbjct  81   DGFGVLRRLRADGIDAPALFLTARDSLQDKIAGLTLGGDDYVTKPFSLEEVVARLRVILR  140

Query  122  QA----EPAAKSEAVAQDIAARVA-----------SLSPRERQVMEGLIAGLSNK-----  161
            +A        +      DI                SLSP E  ++   +
Sbjct  141  RAGKGNKEPRNVRLTFADIELDEETHEVWKAGQPVSLSPTEFTLLRYFVINAGTVLSKPK  200

Query  162  ----LIAREYDISPRTIEVYRANVMTKMQANSLSELVRLAMRAGML  203
            +   ++          +E Y + +  K+       L+      G +
Sbjct  201  ILDHVWRYDFGGDVNVVESYVSYLRRKIDTGEK-RLLHTLRGVGYV  245
</pre></td>
</tr>
<tr>
<td>3Q9S</td>
<td>2e-38</td>
<td><pre>
Query  6    HIYVIDDDAAMRDSLNFLLDSAGFGVTLFDDAQAFLDALPGLSFGCVVSDVRMPGLDGIE  65
             I VI+DD  + + L   L  AG+ V   D A   L          ++ D+ +P  DG +
Sbjct  39   RILVIEDDHDIANVLRXDLTDAGYVVDHADSAXNGLIKAREDHPDLILLDLGLPDFDGGD  98

Query  66   LLKRMKAQQSPFPILIMTGHGDVPLAVEAMKLGAVDFLEKPFEDDRLTAMIESAIRQAEP  125
            +++R++   S  PI+++T    V   V  + LGA D+L KPF  D L A ++  +RQ
Sbjct  99   VVQRLRKN-SALPIIVLTARDTVEEKVRLLGLGADDYLIKPFHPDELLARVKVQLRQRTS  157

Query  126  AAKSEAVAQDIAAR--------VASLSPRERQVMEGLIA---------GLSNKLIAREYD  168
             + S      +              LSP+E  ++  LI           +  ++
Sbjct  158  ESLSXGDLTLDPQKRLVTYKGEELRLSPKEFDILALLIRQPGRVYSRQEIGQEIWQGRLP  217

Query  169  ISPRTIEVYRANVMTKMQANSLSELVRLAMR  199
                 ++V+ AN+  K++      L+R
Sbjct  218  EGSNVVDVHXANLRAKLRDLDGYGLLRTVRG  248
</pre></td>
</tr>
</table>

| | | |
|---|---|---|
| 4KNY | 2e-38 | Query 3 TKGHIYVIDDDAAMRDSLNFLLDSAGFGVTLFDDAQAFLDALPGLSFGCVVSDVRMPGLD 62<br> ++ +++ D+ A+R L L+ G V + Q L ++ D+ +P D<br>Sbjct 2 AMANVLIVEDEQAIRRFLRTALEGDGMRVFEAETLQRGLLEAATRKPDLIILDLGLPDGD 61<br><br>Query 63 GIELLKRMKAQQSPFPILIMTGHGDVPLAVEAMKLGAVDFLEKPFEDDRLTAMIESAIRQ 122<br> GIE ++ ++ S P+++++ + + A+ GA D+L KPF L A + A+R+<br>Sbjct 62 GIEFIRDLRQ-WSAVPVIVLSARSEESDKIAALDAGADDYLSKPFGIGELQARLRVALRR 120<br><br>Query 123 A------EPAAKSEAVAQDIAAR-------VASLSPRERQVMEGLIAGLSNKLIAREYD- 168<br> +P K V D+AAR L+P E +++ L+ L R+<br>Sbjct 121 HSATTAPDPLVKFSDVTVDLAARVIHRGEEEVHLTPIEFRLLAVLLNNAGKVLTQRQLLN 180<br><br>Query 169 --------ISPRTIEVYRANVMTKMQANSLSELVRLAMRAGM 202<br> + +Y ++ K++ + G+<br>Sbjct 181 QVWGPNAVEHSHYLRIYMGHLRQKLE-QDPARPRHFITETGI 221 |
| 4S04 | 4e-38 | Query 5 GHIYVIDDDAAMRDSLNFLLDSAGFGVTLFDDAQAFLDALPGLSFGCVVSDVRMPGLDGI 64<br> I VI+DDA + L + S G+ A +L + +V D+ +P DG+<br>Sbjct 1 MKILVIEDDALLLQGLILAMQSEGYVCDGVSTAHEAALSLASNHYSLIVLDLGLPDEDGL 60<br><br>Query 65 ELLKRMKAQQSPFPILIMTGHGDVPLAVEAMKLGAVDFLEKPFEDDRLTAMIESAIRQAE 124<br> L RM+ ++ P+LI+T + + + GA D+L KPF + L A I + +R+<br>Sbjct 61 HFLSRMRREKMTQPVLILTARDTLEDRISGLDTGADDYLVKPFALEELNARIRALLRRHN 120<br><br>Query 125 PAAKSEAVAQDIAARVA-----------SLSPRERQVMEGLIAGLS---------NKLIA 164<br> +E ++ V L+P+E ++ L+ N + +<br>Sbjct 121 NQGDNEISVGNLRLNVTRRLVWLGETALDLTPKEYALLSRLMMKAGSPVHREILYNDIYS 180<br><br>Query 165 REYDISPRTIEVYRANVMTKMQANSLSELVR 195<br> + + + T+EV+ N+ K+ S VR<br>Sbjct 181 GDNEPATNTLEVHIHNLREKIG-KSRIRTVR 210 |

| | | |
|---|---|---|
| 2OQR | 4e-38 | Query 2 TTKGHIYVIDDDAAMRDSLNFLLDSAGFGVTLFDDAQAFLDALPGLSFGCVVSDVRMPGL 61<br>+ +++D+ ++ D L FLL GF T+ D A L V+ D+ +PG+<br>Sbjct 2 AMATSVLIVEDEESLADPLAFLLRKEGFEATVVTDGPAALAEFDRAGADIVLLDLMLPGM 61<br><br>Query 62 DGIELLKRMKAQQSPFPILIMTGHGDVPLAVEAMKLGAVDFLEKPFEDDRLTAMIESAIR 121<br>G ++ K+++A+ S P++++T V ++LGA D++ KP+ L A I + +R<br>Sbjct 62 SGTDVCKQLRAR-SSVPVIMVTARDSEIDKVVGLELGADDYVTKPYSARELIARIRAVLR 120<br><br>Query 122 QA--EPAAKSEAVAQDIAARVA-------------SLSPRERQVMEGLIAGLSNKLIARE 166<br>+ + + S+ V + R+ +L +E ++E L+ L +<br>Sbjct 121 RGGDDDSEMSDGVLESGPVRMDVERHVVSVNGDTITLPLKEFDLLEYLMRNSGRVLTRGQ 180<br><br>Query 167 Y--------DISP-RTIEVYRANVMTKMQAN--SLSELVRLAMRAGM 202<br>+ +T++V+ + +K++A+ + LV G<br>Sbjct 181 LIDRVWGADYVGDTKTLDVHVKRLRSKIEADPANPVHLVT-VRGLGY 226 |
| 5F64 | 4e-38 | Query 2 TTKGHIYVIDDDAAMRDSLNFLLDSAGFGVT-LFDDAQAFLDALPGLSFGCVVSDVRMPG 60<br>+ + +IDD ++LL + + + + + L V+ DV +PG<br>Sbjct 1 SNAMNAIIIDDHPLAIAAIRNLLIKNDIEILAELTEGGSAVQRVETLKPDIVIIDVDIPG 60<br><br>Query 61 LDGIELLKRMKAQQSPFPILIMTGHGDVPLAVEAMKLGAVDFLEKPFEDDRLTAMIESAI 120<br>++GI++L+ ++ +Q I+I++ D GA F+ K + + A IE+A<br>Sbjct 61 VNGIQVLETLRKRQYSGIIIIVSAKNDHFYGKHCADAGANGFVSKKEGMNNIIAAIEAAK 120<br><br>Query 121 RQ--AEPAAKSEAVAQDIA--ARVASLSPRERQVMEGLIAGLSNKLIAREYDISPRTIEV 176<br>P + + V + ++SLS +E VM ++ G N IA + IS +T+<br>Sbjct 121 NGYCYFPFSLNRFVGSLTSDQQKLDSLSKQEISVMRYILDGKDNNDIAEKMFISNKTVST 180<br><br>Query 177 YRANVMTKMQANSLSELVRLAMRAGM 202<br>Y++ +M K++ SL +L A R +<br>Sbjct 181 YKSRLMEKLECKSLMDLYTFAQRNKI 206 |

| | | |
|---|---|---|
| 4B09 | 6e-38 | Query 3 TKGHIYVIDDDAAMRDSLNFLLDSAGFGVTLFDDAQAFLDALPGLSFGCVVSDVRMPGLD 62<br> I +++D+ + L L +A + TL L + ++ D+ +PG D<br>Sbjct 9 NTPRILIVEDEPKLGQLLIDYLRAASYAPTLISHGDQVLPYVRQTPPDLILLDLMLPGTD 68<br><br>Query 63 GIELLKRMKAQQSPFPILIMTGHGDVPLAVEAMKLGAVDFLEKPFEDDRLTAMIESAIRQ 122<br> G+ L + ++ + S PI+++T + + +++GA D++ KP+ + A +++ +R+<br>Sbjct 69 GLMLXREIR-RFSDIPIVMVTAKIEEIDRLLGLEIGADDYIXKPYSPREVVARVKTILRR 127<br><br>Query 123 AEPAAK-------------SEAVAQDIAARVASLSPRERQVMEGLIAGLSNKLIAREYD- 168<br> +P + ++ L+P E ++++ L +<br>Sbjct 128 CKPQRELQQQDAESPLIIDEGRFQASWRGKMLDLTPAEFRLLKTLSHEPGKVFSREQLLN 187<br><br>Query 169 --------ISPRTIEVYRANVMTKMQANSLSELVRLAM 198<br> ++ RTI+ + N+ K+++ +A+<br>Sbjct 188 HLYDDYRVVTDRTIDSHIKNLRRKLESLDAEQSFIRAV 225 |
| 4YN8 | 5e-37 | Query 5 GHIYVIDDDAAMRDSLNFLLDS-AGFGVT-LFDDAQAFLDALPGLSFGCVVSDVRMPGLD 62<br> + +IDD +R L +LDS V D + VV+D++MPG D<br>Sbjct 6 IRVMLIDDHPVVRAGLRSILDSFDDITVVAEASDGSN----INTKGIDVVVTDIQMPGTD 61<br><br>Query 63 GIELLKRMKAQQSPFPILIMTGHGDVPLAVEAMKLGAVDFLEKPFEDDRLTAMIESAI-- 120<br> GI L + + A P+LI+T + + A++ GA+ +L K + L + +<br>Sbjct 62 GITLTRAL-ANAGGPPVLILTTYDTEADILAAVEAGAMGYLLKDAPESALHDAVVATFEG 120<br><br>Query 121 RQAEPAAKSEAVAQDIAARVASLSPRERQVMEGLIAGLSNKLIAREYDISPRTIEVYRAN 180<br> R+ + A+ Q ++ +LS RE ++++ L GLSN+ +A + IS T++ + +<br>Sbjct 121 RRTLAPEVANALMQRVSKPRQALSAREIEILQNLEQGLSNRQLAAKLFISEATVKTHLVH 180<br><br>Query 181 VMTKMQANSLSELVRLAMRAGML 203<br> + +K+ ++ + + A + ++<br>Sbjct 181 IYSKLGVDNRTAAITAARQQRLI 203 |

| | | |
|---|---|---|
| 1P2F | 5e-36 | Query 6 HIYVIDDDAAMRDSLNFLLDSAGFGVTLFDDAQAFLDALPGLSFGCVVSDVRMPGLDGIE 65<br>I V+DDD + ++ L G V F + FL+ +F VV DV +P G E<br>Sbjct 4 KIAVVDDDKNILKKVSEKLQQLG-RVKTFLTGEDFLN--DEEAFHVVVLDVXLPDYSGYE 60<br><br>Query 66 LLKRMKAQQSPFPILIMTGHGDVPLAVEAMKLGAVDFLEKPFEDDRLTAMIESAIRQAEP 125<br>+ + +K + ++++T D ++ + GA D++ KPF + L A ++ + + +<br>Sbjct 61 ICRXIKETRPETWVILLTLLSDDESVLKGFEAGADDYVTKPFNPEILLARVKRFLEREKK 120<br><br>Query 126 A--------AKSEAVAQDIAARVASLSPRERQVMEGLIAGLSNKLIAREY-------DIS 170<br>+ + + L +E +++ L + + +S<br>Sbjct 121 GLYDFGDLKIDATGFTVFLKGKRIHLPKKEFEILLFLAENAGKVVTREKLLETFWEDPVS 180<br><br>Query 171 PRTIEVYRANVMTKMQANS 189<br>PR ++ + ++ +<br>Sbjct 181 PRVVDTVIKRIRKAIEDDP 199 |
| 1YS6 | 2e-35 | Query 18 DSLNFLLDSAGFGVTLFDDAQAFLDALPGLSFGCVVSDVRMPGLDGIELLKRMKAQQSPF 77<br>SL L +GF V D L + +V D+ MP LDG+ ++ ++A +<br>Sbjct 21 ASLERGLRLSGFEVATAVDGAEALRSATENRPDAIVLDINMPVLDGVSVVTALRAMDNDV 80<br><br>Query 78 PILIMTGHGDVPLAVEAMKLGAVDFLEKPFEDDRLTAMIESAIRQAEPAAKSEAVAQDIA 137<br>P+ +++ V V ++ GA D+L KPF L A +++ +R+ A S + +<br>Sbjct 81 PVCVLSARSSVDDRVAGLEAGADDYLVKPFVLAELVARVKALLRRRGSTATSSSETITVG 140<br><br>Query 138 ARVASLSPRERQVMEGLIAGLSNKLIAREYDISPRTIEVYRANVMTKMQANSLSELVRLA 197<br>+ R +V G+ L RE+D+ E ++ V+++ Q L A<br>Sbjct 141 PLEVDIPGRRARV-----NGVDVDLTKREFDLLAVLAE-HKTAVLSRAQLLELVWGYDFA 194<br><br>Query 198 MRAGMLN 204<br>+++<br>Sbjct 195 ADTNVVD 201 |

*Table S2. PSI-BLAST alignments of FixJ$_{PDB}$ (PDB ID 5xso, HTH$_4$ fold) with sequences of experimentally determined isolated wH folds (upper table). PSI-BLAST alignments of KdpEJ$_{PDB}$ (PDB ID 4kfc, wH fold) with sequences of experimentally determined isolated HTH$_4$ folds (lower table). In both tables, regions with differing secondary structures are bold.*

| Query<br>Subject | Aligned query seq<br>Aligned subject seq | E-value |
|---|---|---|
| 5xso_A | MIESAIRQAEPAAKSE**AVAQDI**AARVASLSPRERQVMEGLIAG----<br>-----LSNKLIAREYDISPRTIEVYRANVMTKMQANSLSEL**VRLAMRAGM** | 6.65e-04 |
| 1gxq_A | AVEEVIEMQGLSLDPT**SHRVMA**GEEPLEMGPTEFKLLHFFMTHPERV<br>YSREQLLNHVWGTNVYVEDRTVDVHIRRLRKALEPGGHDRM**VQTVRGTGY** | |
| 5xso_A | MIESAIRQAEPAAKSE**AVAQDI**AARVASLSPRERQVMEGLIAG----<br>-----LSNKLIAREYDISPRTIEVYRANVMTKMQANSLSEL**VRLAMRAGM** | 7.81e-04 |
| 1qqi_A | AVEEVIEMQGLSLDPT**SHRVMA**GEEPLEMGPTEFKLLHFFMTHPERV<br>YSREQLLNHVWGTNVYVEDRTVDVHIRRLRKALEPGGHDRM**VQTVRGTGY** | |
| 5xso_A | LSPRERQVMEGLIAGLSNKLIAREYDISPRTIEVYRANVMTKMQANS-**LSELVRLAMRAGM** | 0.001 |
| 4qwq_A | LASRENEVISK--SELLEKVWGYDYYEDANTVNVHIHRIREKLEKESF**TTYTITTVWGLGY** | |
| 5xso_A | LSPRERQVMEGLIAGLSNKLIAREYDISPRTIEVYRANVMTKMQANS-**LSELVRLAMRAGM** | 0.002 |
| 4u88_B | LASRENEVISK--SELLEKVWGYDYYEDANTVNVHIHRIREKLEKESF**TTYTITTVWGLGY** | |
| 5xso_A | **ESAIRQAEPAAKSEAVAQDIAARVASL**SPRERQVMEGLIAGLS-------<br>--NKLIAREYDISPRTIEVYRANVMTKMQANSLSEL**VRLAMRAGML** | 0.005 |
| 6kyx_A | **KDIIDVNGITIDKNAFKVTVNGAEIEL**TKTEYDLLYLLAENKNHVMQREQ<br>ILNHVWGYNSEVETNVVDVYIRYLRNKLKPYDRDKM**IETVRGVGYV** | |

| Query<br>Subject | Aligned query seq<br>Aligned subject seq | E-value |
|---|---|---|
| 4kfc_A | **VDLAARVIHRGEEEVHL**TPIEFRLLAVLLNNAGKVLTQRQLLNQVWGPNAVEHSHYLRIYMGHLRQKLEQDPARPRHF | 2.64e-04 |
| 2rnj_A | **VPRGSHMKKRAELYEML**TEREMEILLLIA----KGYSNQEIASA-----SHITIKTVKTHVSNILSKLEVQDRTQAVI |  |
| 4kfc_A | **VIHRGEEEVHL**TPIEFRLLAVLLNNAGKVLTQRQLLNQVWGPNAVEHSHYLRIYMGHLRQKLEQDPARPRHF | 0.004 |
| 7ve4_A | **MKKRAELYEML**TEREMEILLLIA----KGYSNQEIASA-----SHITIKTVKTHVSNILSKLEVQDRTQAVI |  |
| 4kfc_A | **HL**TPIEFRLLAVLLNNAGKVLTQRQLLNQVWGPNAVEHSHYLRIYMGHLRQKLEQDPARPRHF | 0.005 |
| 4wsz_B | **DL**TNREHEILMLIAQGK----SNQEIADELF-----ITLKTVKTHVSNILAKLDVDDRTQAAI |  |

*Table S3 The transitive homology path of 5 sequences between FixJ$_{PDB}$ and KdpE$_{PDB}$. A diagram of the path is shown below.*

| Sequence A | Sequence B | Pairwise identity |
|---|---|---|
| FixJ | WP_007679868 | 45% |
| WP_007679868 | PWU22963 | 38% |
| PWU22963 | TME68356 | 55% |
| TME68356 | WP_021443528 | 38% |
| WP_021443528 | HEK56308 | 43% |
| HEK56308 | KdpE | 48% |

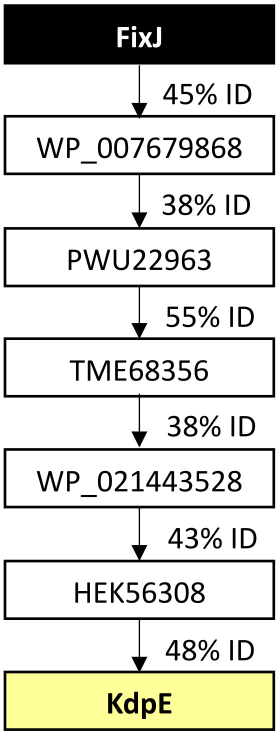

*Table S4. The 12 sequences in the branch containing the bridge sequences; the sequence also found in the transitive homology path (Table S3; also called the "bridge sequence") is highlighted in yellow.*

| GenBank Accession Number | Sequence |
|---|---|
| TMD58691.1 | MGESGPRAGVLLVVEDHVGVRSLVSLVLTAAGYVVREAASGEEAIEFARKEQPLLVLLDV<br>RLPGISGYEVCGWLRDRFHDSVPVIFLSGERTEAFDRAAGLMLGADDYLVKPFSNEELVA<br>RVRGLLRRTLPAPRARGVGLTARELEVLRLLAGGLIQNDIAGHLLISTKTVGTHIEHILM<br>KLGVQSRAQAVALAYRDNLIEPNAPLPIAPSASGPATLSNGAHGDASVSFRTTTAPKPAR |
| TMC70452.1 | SLITAGEPKRSADPNLRAHVQSPLSRAPCAASGVIAGRWDVSGRDGQRRHAVLVVEDDIE<br>VRSLITDVLTHAGYIVRGVQSGEEAIDSVRQEDPILVLVDVHLPGMSGYEVCGWLRSRYR<br>DSVPVMFISGERVEAFDRAAGLMLGADDYMSKPFSTEELVARVRGLLRRTVPPGQSLDTK<br>LTVRELEVLRLLAGGLGQKEISGHLSISAKTVGTHIEHILMKLGVQSRTQAVALAYRERL<br>IDAEDEGESEVRGPTSAKPTS |
| TME68356.1 | MARTVLAWAPTRRSLKRPGTRHGGTRVASSSPRGTILLVDDDPDIRLLLKTVLCNAGFAT<br>NEAVSGEAAVETMRREQPLLVVLDVRLPGVSGYEVCRWLRERFRDIVPIIFMSAERKESF<br>DRAAGLMLGADDYMMKPFSVDELIARIRGLLRRTVPSPRLLAESLTARELEVLRLLAGGL<br>TQADIANQLLISGKTVGTHIEHILMKLDVPSRAQAVAVAYRENLVEPTVAMPTITLTTAS<br>RPSSFGASLLPAATLRMASVAKAAR |
| TMC70453.1 | MSAGGMTVGGPGPRGAILVVEDDADIRSLLSTVLTGAGFAVNTVASGEAAVELMRREQPV<br>LVLLDIRLPGLSGYEVCRWLRERFRDVVPIVFMSAERKESFDRAAGLMLGADDYLMKPFL<br>NEELLARIRGLLRRTVPTPRLLAENLTARELEVLRLLAGGLTQADIANQLLISGKTVGTH<br>IEHILMKLDVPSRAQAVAVAYRENLVEPNGAAPVGPWSTRRVPPTFGSTAVPPATLRLSG<br>QTKSARSG |
| TMB79520.1 | MSELVLVVDDDPNVRGLIVNLLSDYGFETAEASCGEEALRFARERPPDVVLLDVLMPGLS<br>GYEVLRKLKDDFGDSVGVILISGERRESYDRVAGLLVGADDYLPKPFALDELLARVQKLV<br>RGKRRGLPGKRRLTRRELEVLAHLVAGLDNAQIAQRLVLSPRTVTTHVEHILMKLGVHSR<br>TQAIAMAYREGFFEGPSLRVIEGELAAAE |
| OLE96773.1 | MLVVDDDPVICDLVATTLADQGYATRRASDAREALHLIELETPDVVLLDVHLPDLSGYQL<br>CRRLRDTLGDTMGIMLISGERREAFDRAAGLLLGADDYLVKPFVLDELLARVHRMAQRAR<br>PVTLSVAARLTRREAQVLRMLAAGLEQKDIARDLVVAPRTIAKHIEHILLKLGVHSQAQA<br>IALAFRTELAGAVTPHDREEIRVEGT |
| TMC42982.1 | MREGGASMGNPILIVDDDSIIRAAIATILADAGYATREADSGAEALRVARSDAPGLVLLD<br>VNLPGMCGYEVCRLLRDEFGDQFPIVFVSGARTESFDRVAGLLLGANDYISKPFREDELL<br>ARVQSLLLRRHAVASRALASRLTARELQVLRLLSTGLGPDDIARLMVISPKTVGAHVEHI<br>YMKLGVQTRAQAVAVAYRGELLNGETAPTLEPSGNTSR |
| TML48806.1 | MAAATHDGRFVLVVDDDDTFRSLVSEILHRGGYRTRGAATGEEALRVARKKRPSFVLLDV<br>NLPGMSGYDVCRELRAEFGEQLPIVFVSGERTERFDRVAGLRLGADDYIVKPFDPSELLA<br>RVDRFVGRAQAIFSEAATSPFRLTKRELEVLDYLVRGFTPKETARELTISRKTVATHIQN<br>ILTKLDVHSQAQAVAVALQSSLFDFSTRDSESGERSRAHSR |
| TML69799.1 | MTIDCGRGGILVIDSDDEGRAVVSALLADAGYITREAATGREGMSAARRERPDLVLLEVE<br>LRDMTGYEVCRQLRDAYGEGLPVIFVAATRTGSADRIAGLLIGADDYVVKPFAPDELIAR<br>VRRALIRTMSLSRHRNGSRSSYGLTEREREVLRLLSQGLPQKTIARELFISPQTVATHIQ<br>RTLAKLDVHSRAEAVALAHRERLVDDGVGLSAVPEAAP |
| TML69678.1 | MRYNLGAHRRGAQSGERGRVVDCGTVLVVDADDVWRSATAALLTRVGHDTIGAATAADAL<br>ELARRERPSLVVLDVALPDLDGYEVCRELRDVYGDKLPIILVSADRNEPHDCIAALLIGA<br>DDYLAKPYNPGELLARVHRHLTRTNGSNDLTQPKPNTDGSDGFGLSPRELEVLRLLASGL<br>DQAAIAQELVISPKTVSSHIQRILAKLGVHSRAQAVAVAHQEGLVADFAAHAVVFAAPAP<br>SG |
| PWU22963.1 | MRGPILIVDDDPATRELVACTLGSAGYDTLELPTGEQALIAAGEERPALVVLDVKLPGVS<br>GYEVCRQLRERFGEQLPILFVSGERAEAHDRVAGLMIGADDYVTKPFLPDELLARVGRLL<br>VRAKPPAVAADDDLERWAQLTDREREVLNLLAEGLSQDAIAERLYISPKTVATHIQRILA<br>KLGVHSRAAAVSRAYRLGLVSPDFATHMFAFSDA |

*Table S5. Ancestral sequences reconstructed from the consensus phylogenetic tree (Figure S5)*

| *Name* | *Sequence* |
|---|---|
| Anc0 | TATASMLARVLVVDDDPAIRELLADALEREGYRLVVAATGEEALEAAVRGEQRPDLVLLDLALPDLDGF<br>EVCRQLRQQRSNSVPVLFLTGRGEEDDKLRGFRAGADDYVTKPFGPEELLARIRALLRRSRATSPAAAA<br>PSLADAGDDLTLDPARQQATLGLRQPAGLTPREIDLLRLLAQGPGRVVSNREIGEQLGGSELEDADAAR<br>KTVETHVQRLLQKLGADAVSDSRAGAAVQAVAYRLAGSSPSG |
| Anc1 | TSATSPRARILIVDDDPAIRDMLRDALDREGYEVVVAASGEAALEAAVRQEQRPDLVLLDVMLPGMDGF<br>EVCRQLRDQRSNSVPIIFLTGHGDEDDRVAGLRVGADDYVTKPFSPDELLARIRALLRRSRPTRPAAAA<br>PSALRAGDDLSLDPAQQEATLGLQQPAGLTPRETEVLRLLAQGPGRVVSNKEIAEQLGVGEYEDADAAT<br>RTVQVHVRRILKKLGDDGVSHSRAAAVVRGVAYRLAAPSPPS |
| Anc2 | GATASMRARILVVDDDPAIREALRDALERAGYELVTAASGEEALEMAVRNEQQPDLVLLDVMMPGLDGF<br>EVCRQLRQQRSNSVPIIFLTGHGEEDDRVAGLQAGADDYVTKPFSPDELLARIRALLRRSRAASPAAAA<br>PSVLDAGDDLTLDPARQQATLGLQQPAGLTPRETELLRFLAQGPGRVVSNKEIAEQLGISEYEDADAAT<br>RTVQTHVQRLLQKLGDDGVSHSRAAAAVRALAYRLGASSPSS |
| Anc3 | TSATSMMARILIVEDDPAIREMLRDALEREGYEVTTAASGEAALEAAVRQEQRPDLVILDVMLPGMDGF<br>EVCRRLREQSSNSVPIIMLTGRGEEADRVAGLELGADDYVTKPFSPAELLARIRAVLRRSSPAAPAAAP<br>SSVLRFGDDLTLDPARREVTLGAGQPVELTPKEFELLAFLAQRPGRVFSREEILEQVWGPEYEDADAAT<br>RTVEVHISRLRKKLGDNSPSNPRYLLTVRGVGYRFAAPSSES |
| Anc4 | GAAASGRGAILVVDDDPAVRSLIAAILARAGYRLREAATGEEALEAAARGEQRPDLVLLDVRLPGLSGY<br>EVCRQLRDQRGDQLPIIFVSGERESDDRVAGLLIGADDYLTKPFSPDELLARVRRLLRRTHAASPAAAA<br>PSLADAGDDLTLDAPRNDDTLGLSRAAGLTPREREVLRLLAQGPARVLSQKEIAQQLVISPLEDADAAT<br>KTVRTHVQRILAKLGDDGVAHSRAQAVVRALAYREGLVDPDS |
| Anc5 | GGSGPRRGAILVVEDDPDIRSLLSTVLTRAGFRLNEAASGEAAVELMARGEQRPLLVLLDVRLPGLSGY<br>EVCRWLREQRRDVVPIIFMSAERESDDRAAGLMLGADDYLMKPFSNEELLARIRGLLRRRTVPSPAAAA<br>PSLADAGDDLALDSPRNDGTLGLSLAENLTARELEVLRLLAQGPARVLSQADIANQLLISGLEDADAAR<br>KTVHTHVQRILMKLDDDAVAPSRAQAVVRAVAYRENLVEPNA |

### References


1 Eddy, S. R. A new generation of homology search tools based on probabilistic inference. *Genome Inform* **23**, 205-211 (2009).

2 Edgar, R. C. MUSCLE: multiple sequence alignment with high accuracy and high throughput. *Nucleic Acids Res* **32**, 1792-1797, doi:10.1093/nar/gkh340 (2004).

3 Ashkenazy, H. *et al.* ConSurf 2016: an improved methodology to estimate and visualize evolutionary conservation in macromolecules. *Nucleic Acids Res* **44**, W344-350, doi:10.1093/nar/gkw408 (2016).

4 Sievers, F. *et al.* Fast, scalable generation of high-quality protein multiple sequence alignments using Clustal Omega. *Mol Syst Biol* **7**, 539, doi:10.1038/msb.2011.75 (2011).

5 Altschul, S. F. *et al.* Gapped BLAST and PSI-BLAST: a new generation of protein database search programs. *Nucleic Acids Res* **25**, 3389-3402, doi:10.1093/nar/25.17.3389 (1997).
